# Senile plaques in Alzheimer’s disease arise from Aβ and Cathepsin D-enriched amyloidogenic mixtures out of intravascular hemolysis and Charcot-Bouchard aneurysm-related vascular degeneration

**DOI:** 10.1101/2020.05.17.100941

**Authors:** Hualin Fu, Jilong Li, Peng Du, Weilin Jin, Guo Gao, Daxiang Cui

## Abstract

The mechanism governing senile plaque generation in Alzheimer’s disease (AD) remains intensively debated. We analyzed AD brain tissues with histochemistry, immunohistochemistry and fluorescence imaging. We found little co-expression between neural markers and plaque Aβ while abundant co-expression between blood or plasma markers such as HBA, HbA1C, Hemin, ApoE and plaque Aβ. The cores of dense-core plaques were structured with vascular proteins and glial processes at the periphery and blood metabolites in the center. Senile plaques additionally co-localized with a characteristic Hoechst-staining-independent blue autofluorescence, likely also derived from red blood cells. Aβ interacts with hemoglobin in an *in vitro* assay and also at single cell levels in red blood cells *in vivo*, showing as dots, stripes or diffusive patterns with its intensity correlated with coagulation, the elevation of calcium and blue autofluorescence. Interaction between Aβ and ApoE in the blood stream forms vascular amyloid plaques that restrict red blood cell passage. The interaction of Aβ and red blood cells associates with multiple blood and vascular defects besides CAA, including increased perivascular space, microaneurysm, intravascular hemolysis and vascular calcification. Senile plaque formation was intrinsically linked to vascular degeneration as shown by LRP1, ColIV and ACTA2 immunostaining. Aβ staining also overlapped with Cathepsin D expression in intravascular hemolysis, CAA, microaneurysm and senile plaques in AD brain tissues. Microaneurysms with chronic hemolysis were identified as important sites of amyloid formation besides CAA and intravascular hemolysis. In summary, our data suggested that senile plaques arise from Aβ and Cathepsin D-enriched amyloid mixtures out of intravascular hemolysis and Charcot-Bouchard aneurysm-related vascular degeneration. In addition, hemoglobin could be a primary physiological target of Aβ.

## Introduction

Alzheimer’s disease has several important pathological hallmarks such as senile plaques, dystrophic neurites, neurofibillary tangles and cerebral amyloid angiopathy (CAA)^1–4^. Beta amyloid peptide (Aβ) turned out to be a major component in both senile plaques and CAA^5,6^. However, the mechanism governing Aβ generation in senile plaques and the linkage between CAA and senile plaques is still not clear. Many researchers believed that senile plaques are derived from neuronal cells as proposed in amyloid cascade hypothesis^7,8^. On the other hand, there is also strong evidence showing that senile plaques are linked with cerebral microhemorrhage^9–18^. In depth AD mouse model studies based on bone marrow transplantation experiments also indicated that there is a clear role of hematopoietic cell Aβ contributing to senile plaque development^19^. We used immunohistochemistry, fluorescence imaging, TUNEL assay and histochemical staining to examine the neural, vascular or blood Aβ contribution to senile plaque development in AD brain tissues. We found that there is a lack of significant direct neural cell Aβ input to senile plaques. The true direct source of senile plaque Aβ appears to be Aβ-enriched red blood cell (RBC) lysates derived from intravascular hemolysis that come out of cerebral blood vessels through Charcot-Bouchard aneurysm-rupture-related vascular leakage while CAA is an important intermediate stage during this senile plaque formation process.

## MATERIAL AND METHODS

### Tissue sections

AD patient frontal lobe brain paraffin tissue sections were purchased from GeneTex (Irvine, CA, USA). Additionally, AD patient frontal lobe brain paraffin tissue sections were provided by National Human Brain Bank for Development and Function, Chinese Academy of Medical Sciences and Peking Union Medical College, Beijing, China. This study was supported by the Institute of Basic Medical Sciences, Chinese Academy of Medical Sciences, Neuroscience Center, and the China Human Brain Banking Consortium. All procedures involving human subjects are done in accord with the ethical standards of the Committee on Human Experimentation in Shanghai Jiao Tong University and in Xinhua Hospital, and in accord with the Helsinki Declaration of 1975.

### List of antibodies for immunohistochemistry

The following primary antibodies and dilutions have been used in this study: Aβ (Abcam ab201061, 1:200), Aβ/AβPP (CST #2450, 1:200), phos-TAU (Abcam ab151559, 1:200), MAP2 (Proteintech 17490-1-AP, 1:200), GFAP (Abcam ab33922, 1:200), GFAP (Proteintech 60190-1-Ig, 1:100), Iba1 (Abcam ab178847, 1:200), ColIV (Abcam ab236640, 1:200), ACTA2 (Proteintech 23081-1-AP, 1:400), LRP1 (Abcam ab92544, 1:200), ApoE (Abcam ab183597, 1:200), HBA (Abcam ab92492, 1:200), Anti-Glycophorin A antibody (Abcam ab129024, 1:200), Cathepsin D (Proteintech 66534-1-Ig, 1:100), Cathepsin D (Abcam ab75852, 1:200), HbA1c (OkayBio K5a2, 1:100), Hemin (Absolute Antibody, 1D3, 1:100). The following secondary antibodies and dilutions have been used in this study: donkey anti-mouse Alexa-594 secondary antibody (Jackson ImmunoResearch 715-585-150, 1:400), donkey anti-rabbit Alexa-488 secondary antibody (Jackson ImmunoResearch 711-545-152, 1:400), donkey anti-rabbit Alexa-594 secondary antibody (Jackson ImmunoResearch 711-585-152, 1:400), and donkey anti-mouse Alexa-488 secondary antibody (Jackson ImmunoResearch 715-545-150, 1:400).

### Immunohistochemistry

Immunohistochemistry was performed as described^20^. Briefly, paraffin sections were deparaffinized by Xylene, 100% EtOH, 95% EtOH, 75% EtOH, 50% EtOH, and PBS washes. Sections were treated with 10 mM pH6.0 sodium citrate or 10mM pH9.0 Tris-EDTA solution with microwave at high power for 3 times of 4 minutes, with 5 minutes intervals. The sections were allowed to naturally cool down to room temperature. Then, the slides were blocked with TBST 3% BSA solution for 1 hour at room temperature. After blocking, the samples were incubated with primary antibodies at room temperature for 2 hrs followed by 5 washes of TBST. After that, the samples were incubated with fluorescent secondary antibodies overnight at 4 degree. The treated samples were washed again with TBST 5 times the second day and mounted with PBS+50% glycerol supplemented with H33342 nuclear dye (Sigma, B2261, 1 μg/ml) and ready for imaging. To differentiate nuclear staining from blue autofluorescence signal, in some of the IHC experiments, PI (Sigma, P4170, 10 μg/ml) was used as a red nucleus fluorescence dye instead of H33342, which stained the nuclei with blue fluorescence. IHC experiments without primary antibodies were used as negative controls. All experiments have been repeated in order to verify the reproducibility of the results.

### TUNEL assay

TUNEL assay kits (C1089) were purchased from Beyotime Biotechnology (Shanghai, China). TUNEL assay was performed as described in the manufacturer’s user manual. Briefly, paraffin sections were deparaffinized by Xylene, 100% EtOH, 95% EtOH, 75% EtOH, 50% EtOH, and H_2_O washes. Then, the sections were digested with Proteinase K (20 μg/ml) in 10mM TE buffer (pH 7.5) for 15 minutes at room temperature. After Proteinase K digestion, the slides were washed with PBS for 5 times. The slides were then incubated in 50 μl freshly-prepared TUNEL labeling mixture, covered with parafilm and incubated for 1 hour at 37 degree. Afterwards, the slides were washed with PBS for 5 times and mounted with PBS+50% glycerol solution with Hoechst nuclear dye and examined under fluorescent microscopes. To perform double labeling of TUNEL and antibody staining, the incubation time of Proteinase K treatment was shortened and adjusted to allow the simultaneous detection of protein antigens and TUNEL signals.

### Rhodanine staining

The Rhodanine staining kit (BA4346) was purchased from Baso Diagnostics Inc. (Zhuhai, China). Briefly, paraffin sections were first deparaffinized by Xylene, 100% EtOH, 95% EtOH, 75% EtOH, 50% EtOH, and H_2_O washes. After a quick wash in 95% ethanol, the slides were stained in the Rhodanine staining solution (0.01% p-Dimethylaminobenzalrhodanine) at 37 degree for 30 minutes. After staining, the slides were treated with a 95% ethanol quick wash and 5 times wash in water, then mounted with PBS+50% glycerol solution for imaging analysis.

### Alizarin Red staining

The Alizarin Red staining kit C0138 was purchased from Beyotime Biotechnology (Shanghai, China). Briefly, paraffin sections were deparaffinized as described above. Then, the slides were stained in the Alizarin Red staining solution (2%, pH4.2) for 5 minutes at room temperature. After staining, the sections were washed 5 times with water, then mounted with PBS+50% glycerol solution for imaging.

### Slide-based Aβ aggregation assay

Human Aβ40 or Aβ42 peptides (Apeptide Co., Shanghai, China) were first solubilized with DMSO at 1 mg/mL concentration. Human hemoglobin (Hb) (H7379, Sigma) was solubilized in sterile H_2_O at 10 mg/mL concentration. Then, Aβ peptides were diluted into sterile PBS solution at 5μM concentration with or without 1.25 μM hemoglobin. The mixture was incubated at 37 degree up to 3 days. Samples were obtained at Day 0, Day 1, Day 2, and Day 3. 2 μl out of each obtained sample was spotted onto adhesive glass slides and dried up in a 37-degree incubator. The glass slides with samples were fixed with 4% formaldehyde in PBS for 10 minutes at room temperature and washed 3 times with PBS. To test the sensitivity of aggregates to protease digestion, paralleled slides were treated with Proteinase K at 20 μg/mL in 10mM TE (pH 7.5) buffer for 15 minutes at room temperature. The procedure to analyze the Aβ aggregates on the slides using antibodies was the same as in immunohistochemistry.

### Imaging and morphometry analysis

Initially, most images were taken with a Leica DM6000B microscope (Leica Microsystems, Wetzlar, Germany) with A4, L5 and N3 fluorescent filter cubes. The microscope is equipped with 10X eyepiece, and 5X, 10X, 20X, 40X objective lens. During the time of manuscript preparation, slides were re-examined and images were retaken with an Olympus IX71 fluorescent microscope (Olympus Co., Toyoko, Japan), a Keyence BZ-X800 microscope (Keyence Co., Itasca, USA) and a CQ1 confocal fluorescent microscope (Yokogawa, Ishikawa, Japan). To gather more detailed information of tissue sections, DIC or phase-contrast images were also taken along with fluorescent images. When comparing marker immunostaining densities, identical exposure settings for fluorescent channels were used for both control and test groups. Images were then analyzed with Image J software. Mean area density was used as the parameter to define the marker densities. The perimeters of different structures were also measured to simplify the diameter estimation by using the formula: The diameter=The perimeter/3.14. The areas of different structures were also measured with Image J software. The total area intensity was calculated by multiplying the area mean density with the measured area. When using the average neural cell as a reference, the values of other anatomical structures divided the mean value of neural cells to obtain the fold numbers. The folds of changes were then expressed as means with standard deviations.

### Statistics

All data first went through a Shapiro & Wilk normality test using SPSS Statistics 19 software. Two-tailed unpaired T-test was used to compare the means of data with a normal distribution with Excel 2007 software. For data that were not normally distributed, nonparametric Mann/Whitney test was performed to compare the means by using Megastat version 10.2 or with SPSS Statistics 19 software. The p Value threshold for statistical significance is set at 0.05. If p<0.05, then the difference was considered statistically significant. When p<0.001, it was labeled as p<0.001. If p ≧0.001, then the p Value was shown as it was.

## Results

### Neural marker staining rarely overlapped with senile plaque Aβ staining

If senile plaque Aβ comes from neural cells, theoretically, senile plaque Aβ expression should overlap with some neural markers because *in vivo* Aβ does not exist as a single species molecule but rather associates with many other molecules^21^. We checked the expression of neuronal marker MAP2, astroglial marker GFAP, microglial marker Iba1 and a pan-neural marker phos-Tau in AD patient frontal lobe brain tissues. In this study, in order to get a broad image of Aβ-related changes during senile plaque development, “Aβ” refer to all peptides containing the “Aβ” peptide sequence, including the precursor protein AβPP. We found rare staining overlaps between senile plaque Aβ with MAP2, GFAP, Iba1 or phos-Tau(Figure 1). Especially, in the centers of dense-core senile plaques, there is no significant expression of any of the neural markers that we have checked. The data suggested that Aβ from other sources likely form the bulk of Aβ materials in senile plaques, especially in the cores of dense-core senile plaques. In the following analysis, we focused more on studying the structures of the cores of dense-core senile plaques since we consider it is crucial for the understanding of senile plaque pathogenesis.

**Figure 1.**
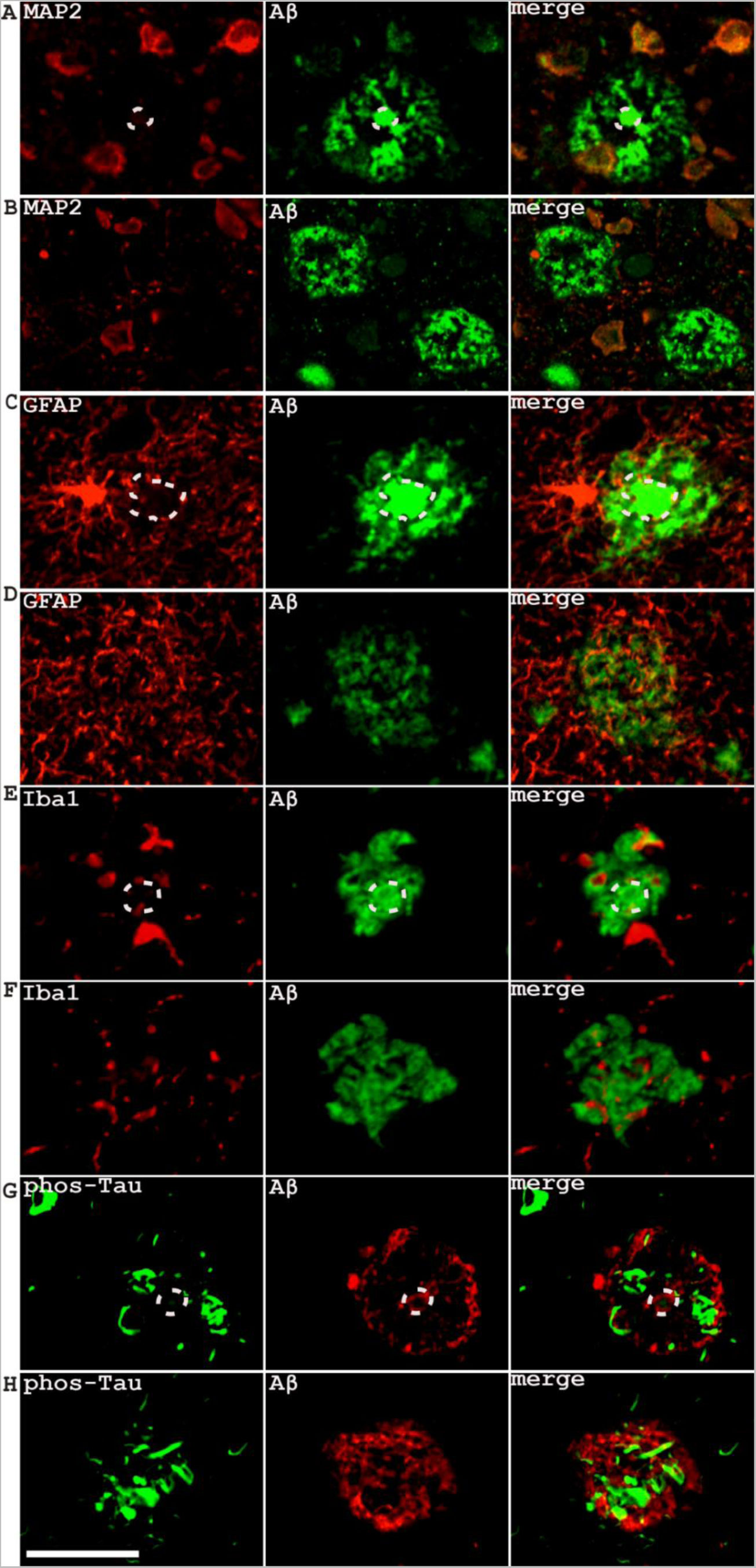
Co-immunostaining of neuronal marker MAP2, glial cell markers GFAP and Iba1, pan-neural marker phos-Tau with Aβ in dense-core or diffusive senile plaques as shown by confocal imaging. (**A**, **B**), MAP2; (**C**, **D**), GFAP; (**E**, **F**), Iba1; (**G**, **H**), phos-Tau. The circled areas indicated the cores of dense-core plaques, which were not stained with any of the neural markers. Scale bar, 50 μm.

### The cores of dense-core senile plaques were characterized by a specific blue autofluorescence unrelated to nuclear staining

Because senile plaque Aβ often shows extracellular localizations, people assumed that Aβ might be derived from dead neurons. However, the direct evidence that links senile plaque Aβ with dying neurons is lacking. Our imaging analysis showed that compact plaques seldom have nuclei associations, agreeing with their predominantly extracellular localizations (Figure 2Aa). However, compact plaques and dense-core senile plaques did not co-localize with Hoechst nuclei stain or TUNEL-labeled nuclei, not supporting the assumption that senile plaque Aβ might come from dying neurons (Figure 2Aa, Figure 2B middle and bottom panel). Although nucleated cells or TUNEL-positive cells could be observed surrounding some diffusive plaques, but these cells had very weak Aβ immunostaining (Figure 2Aa and Figure 2B top panel). There is no sufficient evidence to claim that these cells initiate the diffusive plaque formation process either.

**Figure 2.**
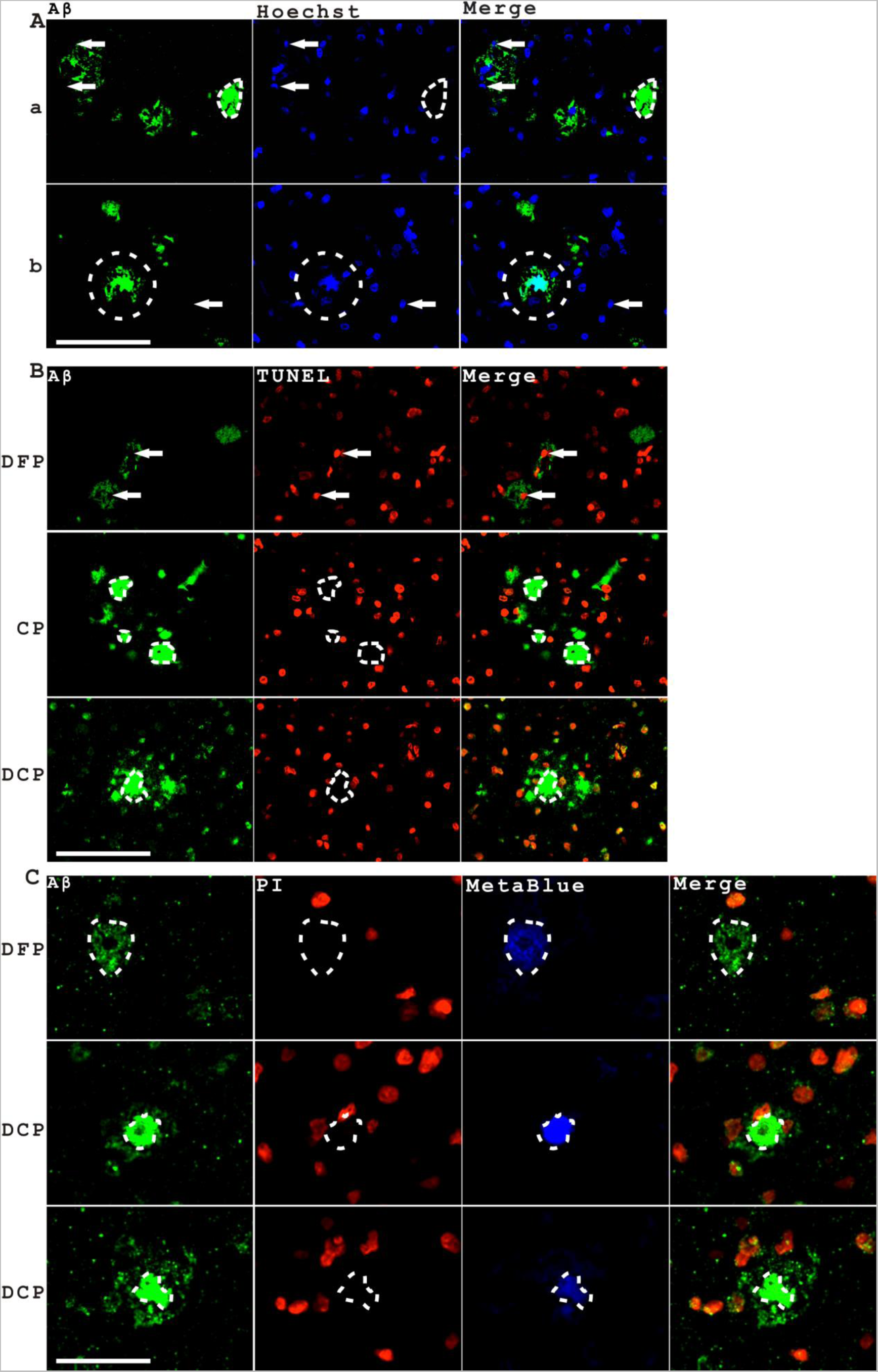
The cores of dense-core senile plaques were characterized by a specific blue autofluorescence, which could not be co-labeled with TUNEL assay or DNA dye PI. **A.** Compact plaques and the cores of dense-core senile plaques were seldom stained with Hoechst staining. Nuclear signals (indicated by arrows) could be observed surrounding diffusive plaques but these nucleated cells did not have significant Aβ staining (**a**). Many compact plaques were negative for Hoechst staining (indicated with dashed lines). However, irregular amorphous blue fluorescent signals were observed in dense-core plaques (DCP) but distinct from nuclear staining patterns (indicated with dash lines in **b**). **B**. Compact plaques and the cores of dense-core senile plaques were seldom labeled with TUNEL staining. TUNEL-labeled cells could be observed surrounding diffusive plaques (DFP) but these TUNEL-labeled cells did not have significant Aβ staining (indicated with arrows in the top panel). Compact plaques were TUNEL negative (indicated by dash lines in the second panel). The cores of dense-core plaques did not contain TUNEL-labeled nuclei (indicated by dash lines in the bottom panel). **C**. The blue fluorescent materials (we named as “MetaBlue” in this manuscript) in the cores of dense-core plaques were not labeled with DNA dye PI. Sometimes diffusive plaques also stained PI-negative (indicated by dash lines in the top panel). The bottom two panels showed the cores of dense-core plaques did not have nuclear staining by PI (indicated by dash lines), however, PI-labeled nuclei can sometimes be seen at the outer rim of dense-core plaques. Scale bar, 100 μm.

Surprisingly, we observed an irregular blue fluorescent staining in the centers of many dense-core plaques, which showed amorphous patterns distinct from densely-stained oval shape nucleus staining (Figure 2Ab). We hypothesized that it was not nuclear staining but instead a type of autofluorescence. To differentiate the blue autofluorescence from the blue nuclei staining, we switched to use a red fluorescence nuclei dye Propidium Iodide (PI). We proved that the amorphous blue fluorescence in the center of dense-core senile plaques is blue autofluorescence, but not nuclei staining (Figure 2C). A strong blue autofluorescence also existed in red blood cells, which provides a first link between red blood cells and senile plaque formation. This blue autofluorescence turned out to be a robust and consistent marker of senile plaques, which associated with senile plaques of different morphologies and sizes even in unstained sections. At the time of preparing this manuscript, we noticed that several earlier studies already recognized senile plaques had characteristic autofluorescence although the true source of this blue autofluorescence is unknown^22,23^. To facilitate the study and discussion on this unique and potentially important substance, we would like to give a temporary name for this material as “MetaBlue” in this manuscript (meaning for a blue metabolite or a mixture of blue metabolites) until the accurate identity of this material is fully revealed.

The lack of nuclear signals nor TUNEL signals in compact plaques and the center of dense core plaques do not support the prevailing hypothesis that senile plaques derive from dying neurons. Although it is still possible that senile plaque Aβ coming from dying neurons with their DNA signals lost very early in the process, there is another possibility that senile plaque Aβ largely come from cells without nuclear DNA at all. Red blood cells and platelets are two main types of cells that do not have nuclear DNA. It is important to study their potential contributions to senile plaque development.

### Significant Aβ staining was detected in some populations of red blood cells with also blue autofluorescence

We already knew that Aβ is expressed in neural cells, in platelets, but we have limited information on the expression of Aβ in red blood cells. In this study, we focused more on the associations between Aβ and red blood cells on pathological sections. Significant Aβ expression in some populations of red blood cells was clearly observed in both CAA vessels and non-CAA vessels (Figure 3A). Aβ associated with some red blood cells at single cell levels, showing different patterns, such as dots, stripes and diffusive patterns (Figure 3B). Red blood cells in AD samples have significant blue autofluorescence similarly as senile plaques as previously stated. The intensity of blue fluorescence in red blood cells was higher than diffusive senile plaques or the diffusive rims of dense-core plaques but lower than CAA or the cores of dense-core plaques (Supplementary Figure 1). Based on one set of confocal images, if using red blood cell average blue autofluorescence as a reference value of 1 (N=107), the relative blue fluorescence intensity in the diffusive plaques, the rims of dense-core plaques, the cores of dense-core plaques and CAA is 0.73±0.37 (N=46), 0.8±0.33 (N=12), 2.04±0.68 (N=12), 1.53±0.76 (N=3) respectively.

**Figure 3.**
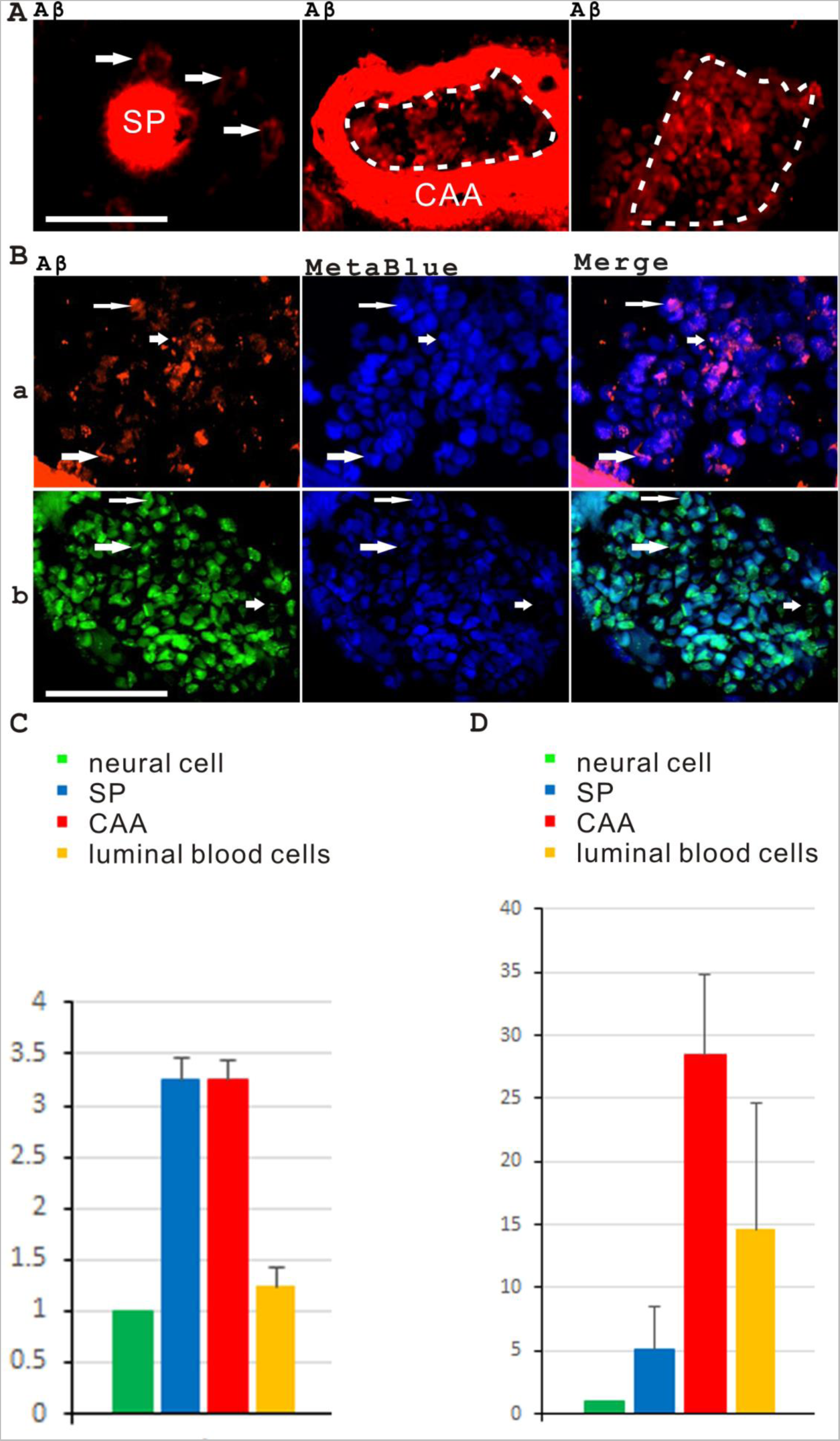
Significant Aβ staining was detected in some populations of red blood cells that also possessed blue autofluorescence. **A**. The left panel showed a senile plaque surrounded by several weakly-stained neural cells. The middle panel showed a CAA vessel with Aβ-positive luminal blood cells. The right panel showed a non-CAA vessel with Aβ-positive luminal blood cells. The arrows indicated the weakly stained neural cells. The dashed lines indicate luminal blood cells. The senile plaques were indicated with “SP” while the CAA blood vessels were labeled as “CAA”. Scale bar, 50 μm. **B.** Aβ associated with red blood cells in dots, stripes or diffusive patterns as shown both with red fluorescence (**a**) and with green fluorescence (**b**). The big and long arrows indicated the stripe patterns, the short arrows indicated dotted staining while the thinner arrows indicated the diffusive staining patterns. Scale bar, 50 μm. **C**. The average Aβ intensity in senile plaques (SP), CAA, Aβ-positive luminal red blood cells is 3.26 (±0.21), 3.26 (±0.18) and 1.24 (±0.19) of the average Aβ intensity in neural cells. **D**. The average area of senile plaques, CAA, luminal regions containing Aβ-positive red blood cells is 5.03 (±3.48), 28.38 (±6.5) and 14.61 (±9.99) of the average area of neural cells. Aβ intensities of luminal Aβ-positive blood cell patches were significantly higher than average neural cells (p=0.018, Mann/Whitney test) while no significant difference was detected between CAA and senile plaques (p=0.943, Mann/Whitney test).

We observed that neural cells express Aβ peptides at very low levels as shown by immunohistochemistry staining. This very low level of Aβ expression in neural cells can be imaged side by side with strong senile plaque Aβ staining with longer exposure settings (Figure 3A). The relative intensity of Aβ expression in 312 neural cells, 125 senile plaques, 9 CAA and 8 Aβ-positive luminal red blood cell patches was quantified in one set of images of Aβ immunostaining. The areas and perimeters of these samples were also measured. By using the microscope scale, the average diameter of senile plaques (calculated from the average perimeter divided by 3.14)is around 38.67 (±15.94) μm. The average area of senile plaques is around 933.6 (±645.15) μm^2^. The average Aβ intensity in senile plaques, CAA, Aβ-positive luminal red blood cells is 3.26 (±0.21), 3.26 (±0.18) and 1.24 (±0.19) times of the average Aβ intensity in neural cells. The average areas of senile plaques, CAA, luminal Aβ-positive red blood cell patches is 5.03 (±3.48), 28.38 (±6.5) and 14.61 (±9.99) times of the average area of a neural cell. The total area density of an average senile plaque is roughly 16.4 times of total area density of an average neural cell. If taking the analysis to the three-dimensional levels, an average senile plaque has around 11.28 times of volume of an average neural cell body and 66.40 times of total Aβ content as in an average neural cell. An average CAA vessel contained 889.90 times Aβ as in an average neural cell, which translated into 13.40 times of Aβ content as in an average senile plaque. An average Aβ positive blood cell patch contained 77.11 times of Aβ as in an average neural cell, which translated into 1.16 times of Aβ as in an average senile plaque. Numerically, these quantitative estimations were close but somewhat different from our earlier calculations^24^, which could be due to different data sets and microscopy methods (regular fluorescent microscope vs. confocal microscope) that were used. The experiment was repeated on different samples and the results showed similarly the extremely low Aβ expression in neural cells comparing to senile plaques. Although Aβ expression level is just one factor that might affect senile plaque formation, the above calculations indicated that, mathematically, besides neural cells, CAA blood vessels and luminal red blood cells could be sources with sufficient amount of Aβ that contribute to senile plaque formation.

### A blue autofluorescence independent of Hoechst staining exists both in the red blood cells and also in the senile plaques

To study the properties of MetaBlue fluorescence, we combined immunohistochemistry staining with fluorescence imaging. First, we verified MetaBlue signals in red blood cells by using red blood cell marker HBA and Glycophorin A antibodies (Figure 4A). Second, we compared the blue fluorescence signals of the same senile plaques with or without Hoechst staining with simultaneous Aβ immunostaining. The results proved that MetaBlue blue autofluorescence was independent of Hoechst staining (Figure 4B). Thirdly, we detected the coexistence of HBA and MetaBlue signals in dense-core plaques and diffusive plaques (Figure 4C). With the senile plaque association of predominantly blood origin MetaBlue and HBA but much less with neural markers such as MAP2, GFAP, Iba1 and phos-TAU, it suggested that senile plaque Aβ might directly come from the blood metabolites instead of neural cells.

**Figure 4.**
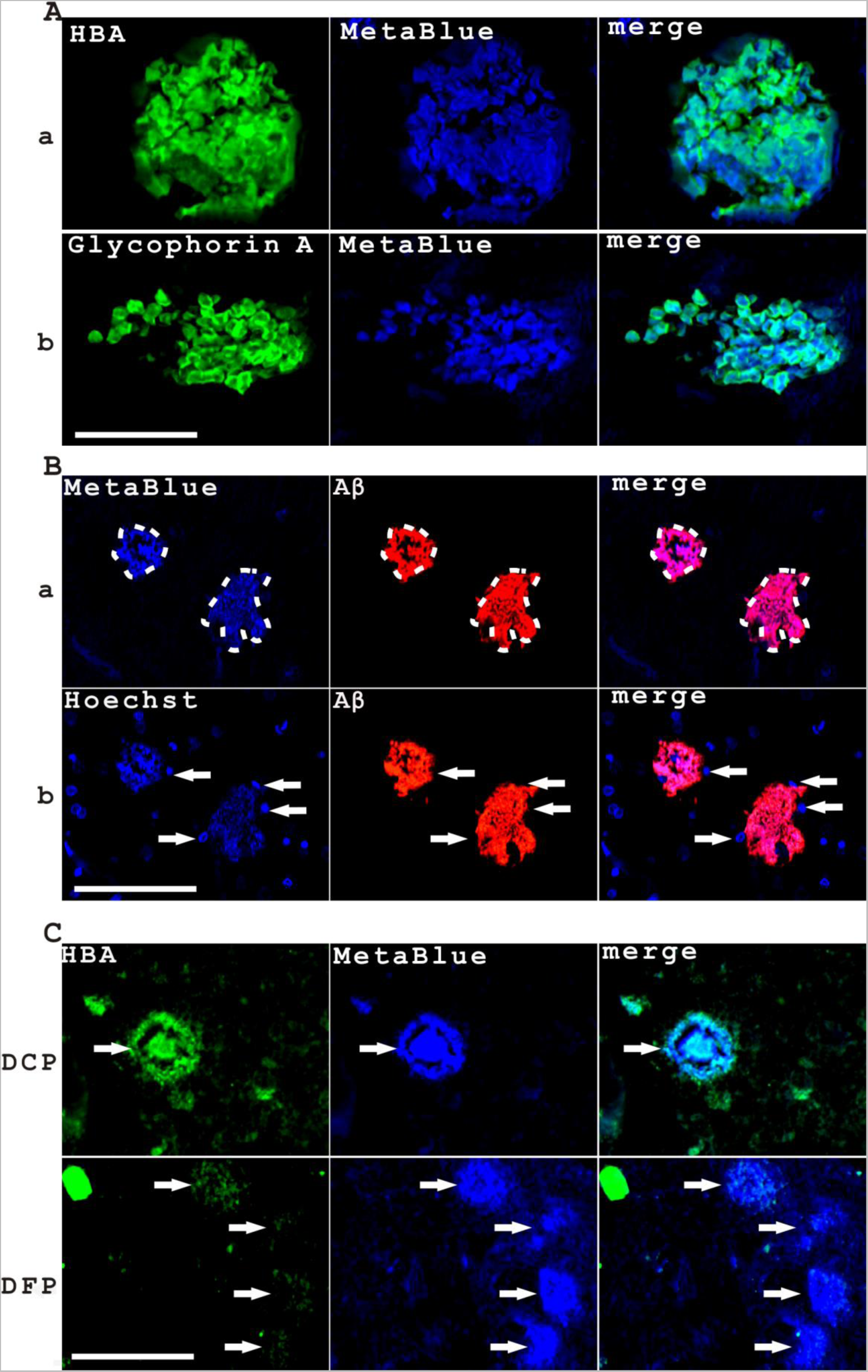
The MetaBlue blue autofluorescence independent of Hoechst staining was present both in the red blood cells and also in the senile plaques. **A**. MetaBlue signals were clearly detected in red blood cells (RBC) labeled either with HBA (**a**) or with Glycophorin A (**b**). Scale bar, 50μm. **B**. MetaBlue blue autofluorescence was independent of Hoechst staining. Hoechst staining (**b**) added some nuclear signals surrounding senile plaques whereas not affecting the pattern of blue autofluorescence (**a**). Scale bar, 100μm. **C**. MetaBlue signals co-stained with HBA staining in dense-core plaques (top), and diffusive plaques (bottom). The arrows pointed to the co-expression of markers in dense-core or diffusive senile plaques. Scale bar, 100 μm.

### The cores of dense-core plaques were intrinsically linked to vascular and blood markers

To understand the nature of the cores of dense-core plaques is essential for elucidating the pathological mechanism of senile plaque formation. We hypothesized that the cores of senile plaques are actually vascular structures with Aβ-associated blood contents, thus, we labeled the dense-core plaques with vascular markers LRP1 and ColIV, astral glial marker GFAP and blood markers HbA1C and Hemin.

The vascular endothelial marker LRP1 labeled the cores of dense-core plaques clearly showing as perfect rings surrounding the dense cores (Figure 5A, B). When the central ring-shape vascular structure disintegrated, a “diffusive” plaque morphology appeared with simultaneous dispersion of LRP1 and Aβ signals (Figure 5C, D). The vascular collagen marker ColIV also labeled the cores of dense-core plaques with degenerating ColIV labeling surrounding the cores of dense core plaques (Figure 5E). The dense cores were also encircled by GFAP labeling, indicating that the dense cores were surrounded by glial processes, which is an important character of cerebral vasculature (Figure 5F). Not surprisingly, we also observed the cores were loaded with blood markers such as HbA1C and Hemin (Figure 5G, H), both highly related to RBC hemoglobin metabolism. In addition, we noticed there was an unusual Aβ distribution pattern in some dense-core plaques. Some dense cores had a strong Aβ signal which forms a ring around the dense cores (Figure 5A, E, F, G and H). The ring shape staining indicated that Aβ in dense cores was asymmetrically distributed, which preferentially enriched at the outer ring of the dense cores corresponding to the luminal surfaces of blood vessels. From static pathological images, it is difficult to know whether Aβ flow out of the dense core or flow into the dense core when the core began to break. However, we fortunately obtained images with dense-cores half broken as judging from the integrity of GFAP-labeled glial foot structures (Supplementary Figure 2). The half core with intact GFAP labeling always had higher Aβ staining than the half core with broken glial foot structures, which suggested that the amyloid formation was pre-formed within the intact core structure. To be noticed, the blood vessels are also internally pressured structures. Considering both concentration gradient and blood pressure gradient, when the core is broken, the correct flow direction should be that Aβ flows out of the core structure rather than Aβ flows into the core. In summary, these results showed that the cores of dense-core plaques were wrapped with vascular markers such as LRP1 and ColIV, and with astral glial marker GFAP at the periphery and also centrally loaded with blood metabolites as labeled by HBA (Figure 4C), HbA1C and Hemin antibodies. In a word, the dense-core senile plaques were derived from vascular structures with blood contents.

**Figure 5.**
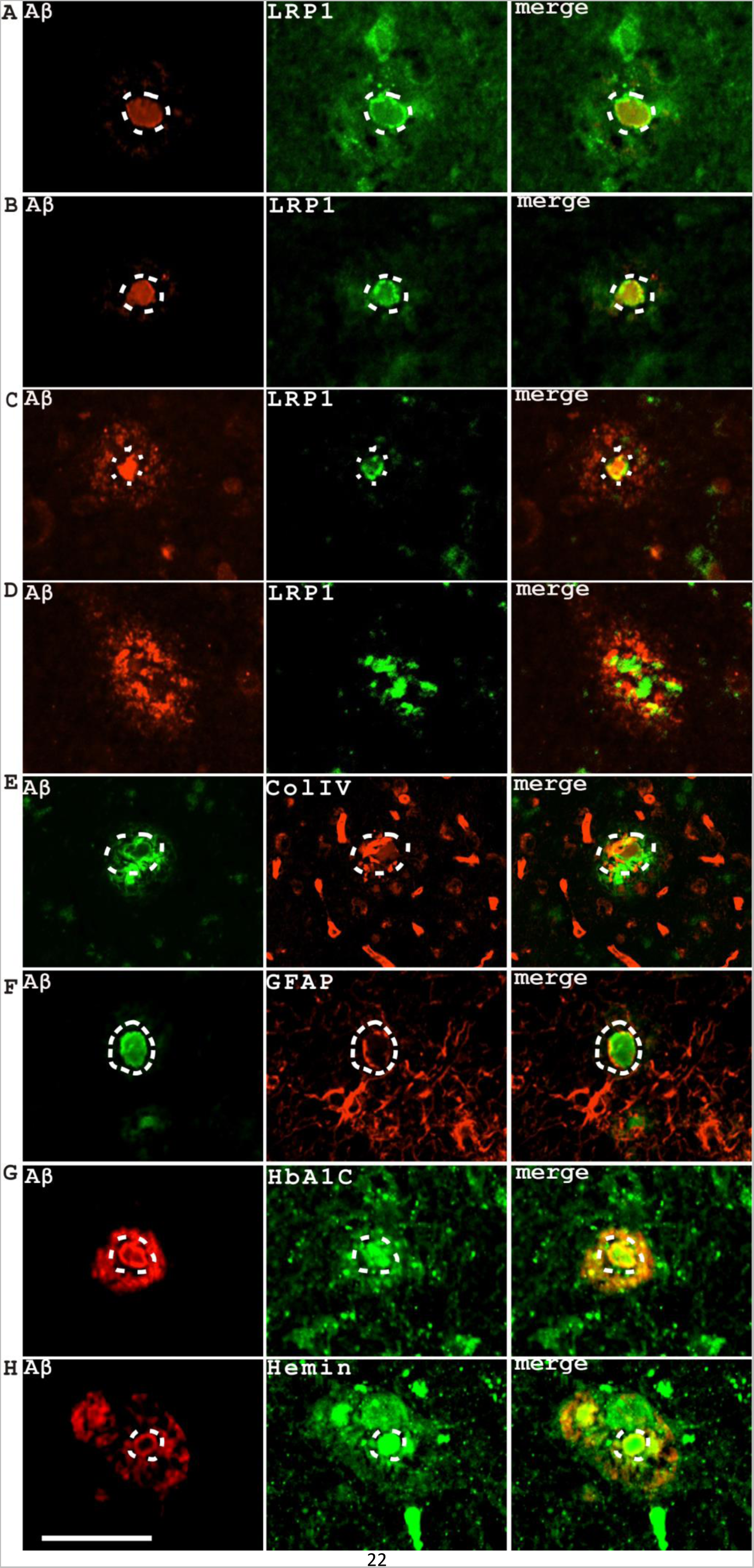
The cores of dense-core plaques were wrapped with vascular, astral glial markers outside and loaded with blood markers inside. (**A, B**): The outer boundary of the cores of dense-core plaques was labeled by vascular endothelium marker LRP1. (**C, D**): The fragmentation of LRP1 labeling was associated with the disintegration of the senile plaque dense cores and the dispersion of Aβ signals. **E.** The outer boundary of the cores of the dense-core plaques was surrounded by ColIV staining, a marker of blood vessels. **F.** The outer boundary of the cores of the dense-core plaques was associated with astral glial marker GFAP, suggesting that the dense cores were surrounded by glial processes. **G.** The center of the cores of dense-core plaques was labeled by red blood cell marker HbA1C (glycated hemoglobin). **H.** The center of the cores of dense-core plaques was labeled by Hemin, an important metabolic derivative of hemoglobin metabolism. The dashed lines indicated the cores of dense-core senile plaques. Scale bars, 100 μm.

### The interaction of ApoE and Aβ in the blood stream contributes to vascular amyloid plaques that restrict blood flow

It is not clear what the physiological consequence of accumulating amyloid beta peptide is in the blood stream. We intended to use other important blood markers besides hemoglobin metabolite markers to study the relation of senile plaque formation and the blood. ApoE is an important plasma protein and a critical risk factor for AD development, which also interacts with Aβ directly^25–27^. Immunostaining experiments showed that ApoE co-expressed with Aβ in dense-core senile plaques, diffusive plaques, and CAA (Figure 6A). Surprisingly, ApoE also co-expressed with Aβ in the lumen of small vessels and capillaries. By using Rhodanine as a red blood cell dye^24^, we observed that the ApoE-Aβ association forms amorphous amyloid structures in the blood vessel lumen that restrict red blood cell passage (Figure 6B). In another word, ApoE-Aβ amyloid complex formation in the blood stream blocks blood flow. We quantified the numbers of vessels or capillaries with strong ApoE and Aβ in the lumens on 5 images (200X magnification) in one set of experiments, we estimated that around 22.9 (±5.9) % of the blood vessels could be affected with blood flow blockage by vascular amyloid plaques in this particular sample. In addition, it was clear that ApoE associated with Aβ in hemolysis, indicated by the reddish color of Rhodanine staining, suggesting the ApoE-Aβ amyloid complexes probably derived from hemolytic products of red blood cells (Figure 6Bd). Since the blockage of blood flow and the formation of vascular plaques are important characters of atherosclerosis, atherosclerosis might be an early driver of Alzheimer’s disease development as already suggested by previous studies^28^.

**Figure 6.**
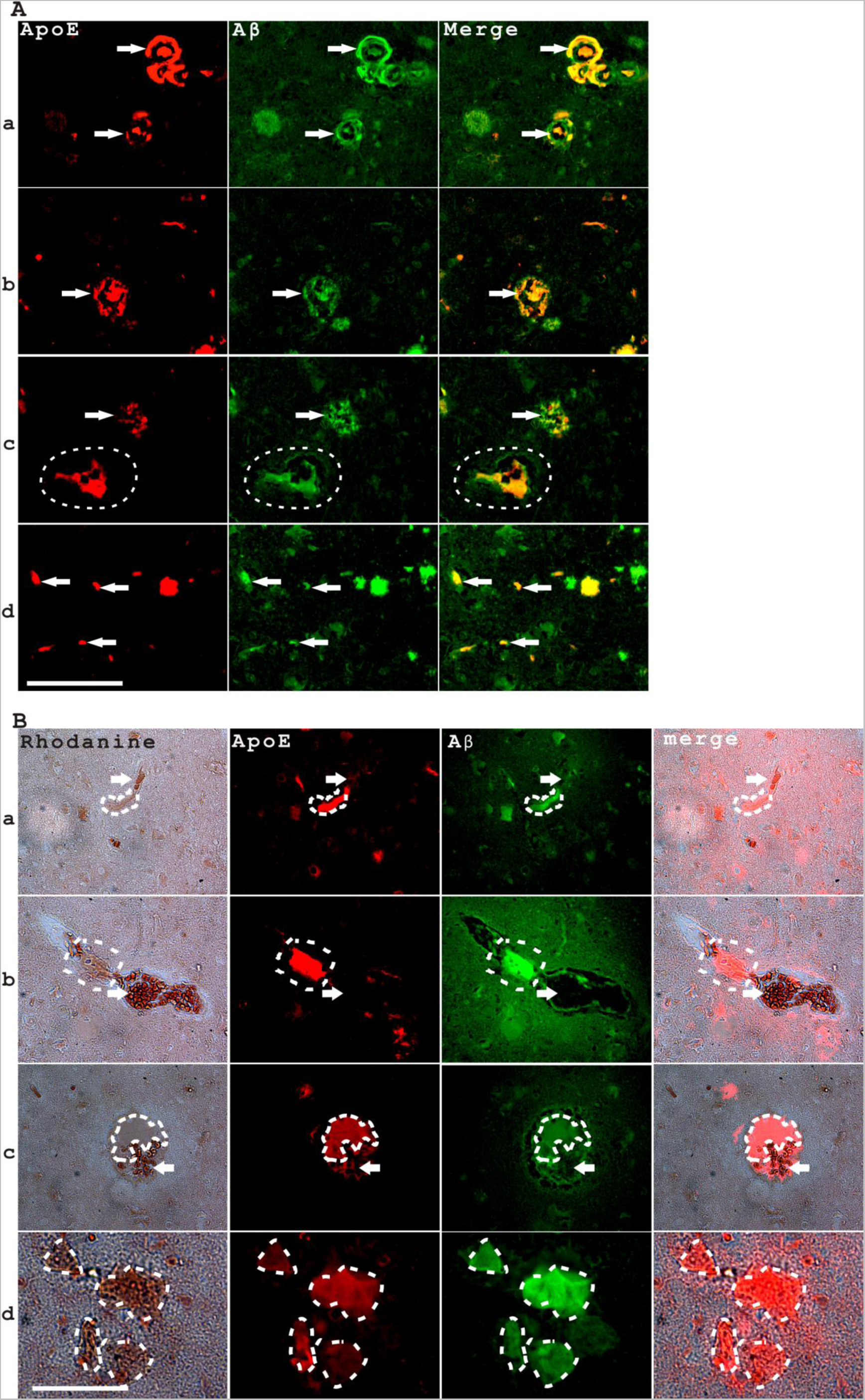
The interaction of ApoE and Aβ in the blood stream contributes to vascular amyloid plaques that restrict blood flow. **A**. ApoE co-expressed with Aβ in CAA, senile plaques and in the blood stream. The top panel showed two CAA vessels with ApoE and Aβ co-immunostaining in both vessels and also in the vessel lumens (indicated with arrows in **a**). The second panel showed a dense-core plaque labeled both with ApoE and Aβ (indicated by arrows in **b**). The third panel showed a diffusive plaque labeled with ApoE and Aβ (indicated by arrows in **c**) and a medium size vessel with luminal ApoE and Aβ staining (indicated by dashed lines). The fourth panel showed simultaneous ApoE and Aβ staining in several capillary size blood vessels (indicated by arrows in **d**). Scale bar, 100 μm. **B**. ApoE associates with Aβ in vascular amyloid plaques that restrict blood cell flow. The top three panels showed that the co-expression of ApoE and Aβ in blood vessels limited blood cell flow in vessels of different sizes, from capillaries to medium size vessels (**a-c**). Dashed lines indicated that the vascular domains of Aβ and ApoE interaction are void of red blood cells (indicated with arrows) labeled deep red with Rhodanine staining. The bottom panel showed that ApoE associates with Aβ in hemolysates (indicated by dashed lines) that can be marked with light red Rhodanine staining (**d**). Scale bar, 100 μm.

### CAA development was intrinsically associated with multiple blood or plasma markers and also blood-related histological stains

CAA is observed in the majority of AD cases^29^. How CAA is developed and how CAA might link to senile plaque formation is not clear. We initially observed MetaBlue and ApoE in CAA while both MetaBlue and ApoE are blood-related markers as previously shown (Figure 7A, B). We hypothesized that CAA development was initiated by the leakage of blood contents into the vascular wall. We stained CAA with more blood-related markers such as hemoglobin related markers and red blood cell specific histological stains. As expected, CAA development was accompanied by the vascular expression of HBA, HbA1c and Hemin (Figure 7C, D, E, F), all of which are the markers of hemoglobin metabolism in red blood cells. CAA association with blood content leakage could be further illustrated by red blood cell histological stains. CAA blood vessels are associated with a pinkish Rhodanine stain which stains red blood cells in bright red (Figure 7G). Further, CAA development is associated with a deep red stain of the vascular wall by Alizarin Red which also stained red blood cells in deep red (Figure 7H). As a control, a non-CAA blood vessel was not stained by Alizarin Red. Blood cells with diffusive Aβ staining were observed in non-CAA blood vessels while in some CAA blood vessels there is a complete loss of red blood cells and the red staining was exclusively residing in the vascular wall. Staining of CAA blood vessels by Alizarin Red, a calcium dye, also suggested that CAA blood vessels are calcified. It is well-known that calcification of blood vessels is associated with vascular aging and atherosclerosis, adding further evidence that atherosclerosis might be an early event in Alzheimer’s disease development.

**Figure 7.**
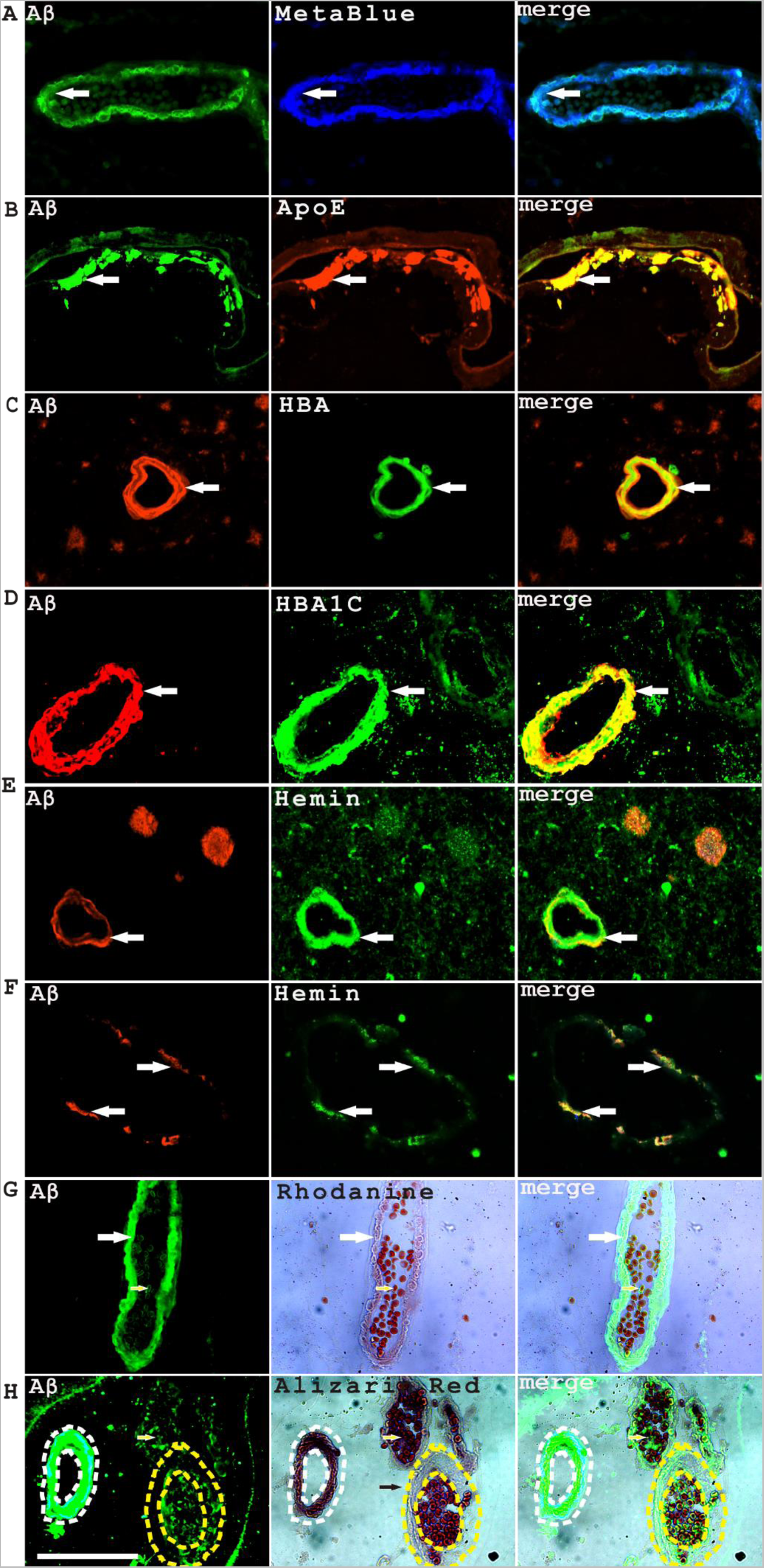
CAA development was intrinsically associated with multiple blood or plasma markers. **A**, MetaBlue; **B**, ApoE; **C**, HBA; **D**, HBA1C; (**E**, **F),** Hemin; **G**, Rhodanine; **H**, Alizarin Red. The arrows indicated the co-expression of Aβ and related markers in CAA. Particularly, the arrows showed a vessel at early stage of CAA development with the accumulation of ApoE and Aβ in the inner layer of the blood vessel and patches of ApoE and Aβ co-expression invading the vascular wall (**B**). Another vessel at early stage of CAA development showed patches of co-expression of Hemin and Aβ scattering in the vascular wall (**F**). Panel **G** showed that CAA vascular wall was labeled by Rhodanine in pink red while red blood cells were labeled bright red by Rhodanine. Panel **H** showed that CAA vascular wall (white dashed lines) and red blood cells (yellow arrows) were both labeled deep red by Alizarin Red while the vascular wall of a non-CAA vessel (yellow dashed lines) was not labeled. Scale bar, 100μm.

In addition to the calcification defect of vascular wall in CAA, around 40% of CAA blood vessels contain no red blood cell at all (Figure 7H, white dashed lines)(38 out of 96 CAA blood vessels counted). A CAA blood vessel with no red blood cells could mean two things: First, the disappearance of preexisting red blood cells might have happened, likely through hemolysis. Second, there is no new red blood cell flowing through this blood vessel, suggesting the upstream of this blood vessel was blocked for blood flow. The interaction of Aβ-ApoE in hemolysis and vascular amyloid plaque formation could explain both aspects.

### Aβ deposition in CAA disrupts blood vessel collagen architecture and decreases ACTA2 expression and vascular cellularity

In addition to the calcification defect in CAA, it is possible that CAA development damages fine vascular wall structures, which also contributes to hemorrhage leakage. We stained CAA blood vessels with collagen IV, a specific marker of vessel wall. We found the collagen layers in the middle layer of blood vessels become disorganized or partially absent (Figure 8A). Instead, the cracks in the collagen layer were filled with Aβ deposition. We further stained the cellular components of blood vessel with an antibody against smooth actin protein ACTA2, a cytoplasmic marker of vascular cells. It is clear that Aβ deposition and ACTA2 staining cells resides in complementary domains in the vessel wall, which suggested that CAA Aβ deposition existed extracellularly, adding further evidence that CAA is developed from blood content leakage (Figure 8B). With increasing Aβ load in CAA, we saw reducing signals of ACTA2. In some CAA vessels, we saw a complete loss of ACTA2 signals indicating a loss of vascular cellularity (Figure 8B, bottom). In addition, we observed that CAA development was accompanying with a gradual loss of nuclear signals by PI staining, further suggesting a loss of cellularity in CAA. Based on the average area intensity quantification of 48 Aβ-and-PI-stained CAA vessels, a weak linear negative relationship between the average PI intensity and Aβ intensity was found (r=-0.287, P=0.048).

**Figure 8.**
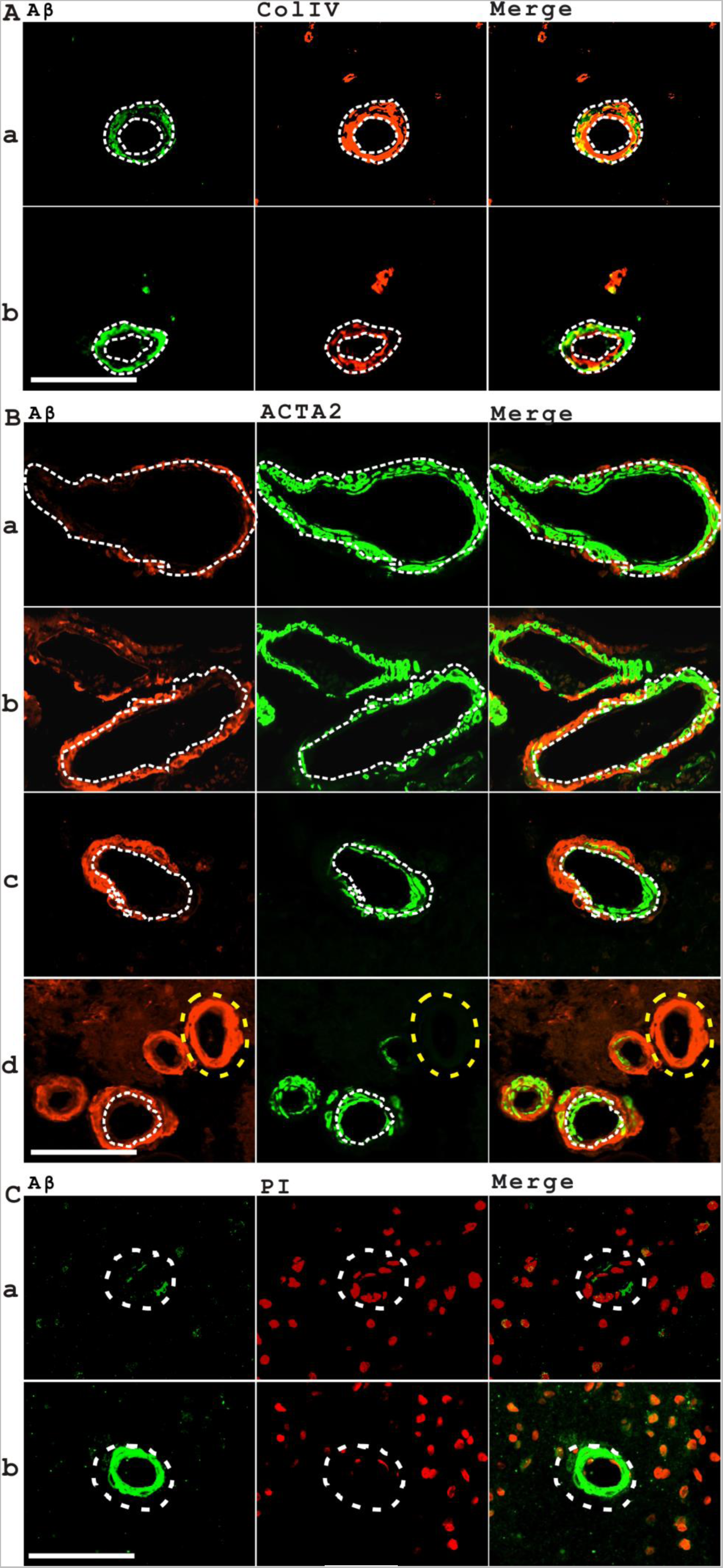
Aβ deposition in CAA disrupts blood vessel collagen matrix architecture and decreases ACTA2 expression with a decrease of cellularity. **A**. Aβ deposition in CAA disrupts blood vessel collagen matrix architecture with two samples shown in **a** and **b** with dashed lines. **B**. Aβ stained complementary locations comparing to ACTA2-expressing cells suggesting that Aβ deposition in CAA might be extracellular (the separation of two domains was indicated by dashed lines in **a** and **b**). The accumulation of Aβ continues to a point of losing all of the smooth-actin expression domains (indicated by yellow dashed lines in **c**). **C**. There is a trend of losing nucleated cells in CAA blood vessels as indicated by PI labeling of cell nucleus. The top panel showed weak Aβ staining in the vascular wall with relatively more nuclei staining (**a**) while the bottom panel showed a CAA blood vessel with strong Aβ staining but less PI-positive nuclear signals (**b**). Scale bars, 100 μm.

### Aβ associates with “increased perivascular space” and aneurysm formation

CAA is a well-known vascular defect that has been linked to abnormal Aβ deposition in Alzheimer’s disease. We are interested to know if there are other blood or vascular phenotypes associating with Aβ deposition. After examining many pathological images, we observed that Aβ associates to at least three additional vascular-related phenotypes: increased perivascular space, aneurysm and intravascular hemolysis. Increased perivascular space was frequently observed surrounding small vessels with the diameter ranging from 20 micrometers to 300 micrometers, with the observed frequency up to 60% on pathological sections (Figure 9B, C). Most of the vessels (around 80%) with increased perivascular space showed the presence of luminal Aβ staining, often associated with red blood cells. Increased perivascular space was also observed surrounding some vessels with massive intravascular hemolysis with strong luminal Aβ staining (Figure 9D). As it has been shown in Figure 6, luminal Aβ staining is often a sign of vascular amyloid plaque formation, which increases flow resistance and restricts blood passage. Microaneurysms with diameters smaller than 100 μm were observed developing from small vessels or capillaries (Figure 9E, F, G, H). All the microaneurysms contained significant amount of Aβ as judged from immunostaining. We noticed cerebral microaneurysms had been studied from the time of Charles Jacques Bouchard with his pioneering study on very small cerebral aneurysm (microaneurysm) named as Charcot-Bouchard aneurysm since 1870s^30^. He hypothesized that saccular or fusiform cerebral aneurysms originating from small vessels with sizes ranging from 250 μm to 400 μm are responsible for the wide spread cerebral microhemorrhage in aging patients.

**Figure 9.**
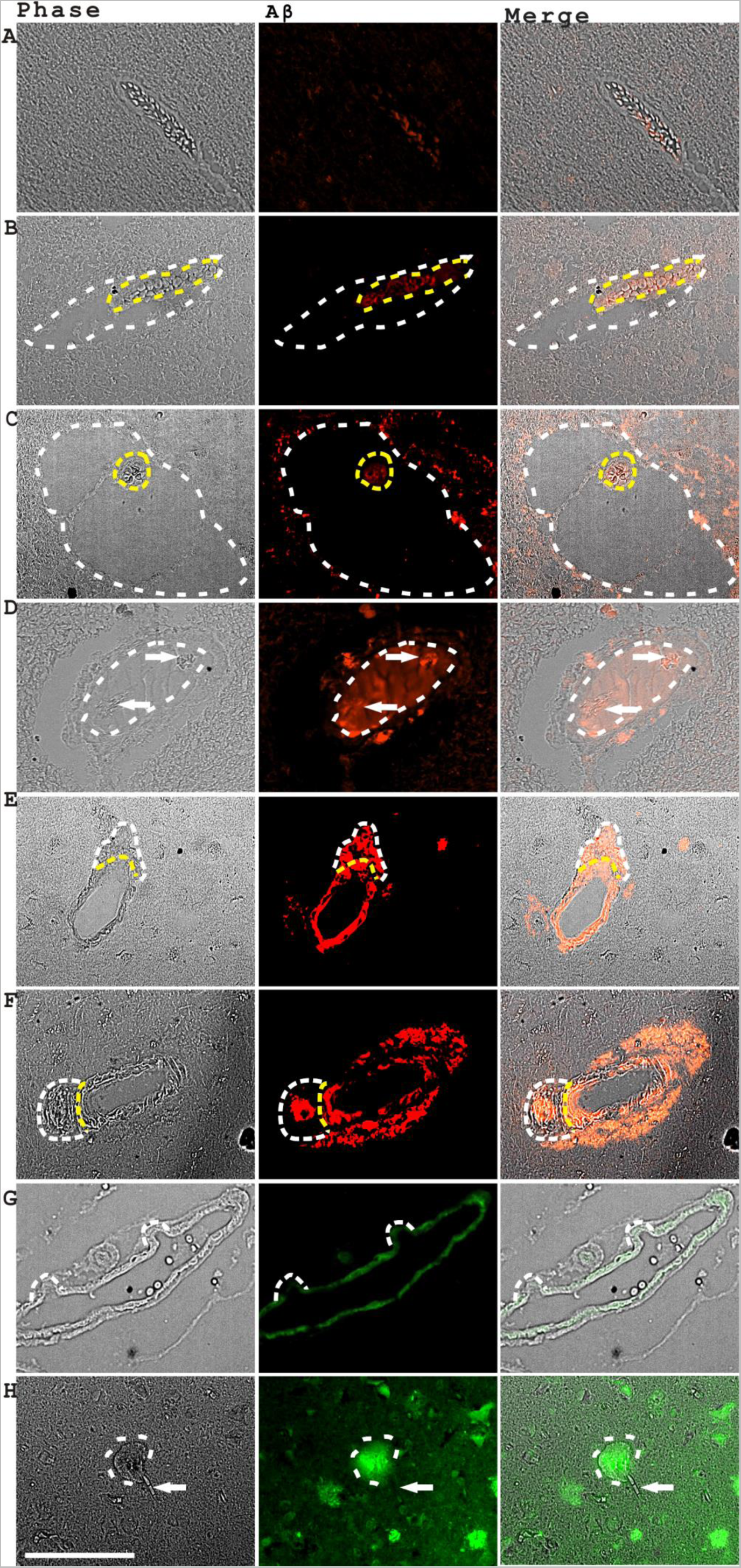
Aβ staining in the blood cells associates with increased perivascular space while Aβ staining in the vascular walls associates with aneurysm formation. **A**. A normal looking blood vessel with smooth vascular wall structure and a few Aβ positive blood cells in the lumen with no “increased perivascular space” phenotype observed. (**B, C**): Two vessels (indicated by dashed lines) with significantly increased perivascular space showed packed Aβ-positive blood cells in the lumen (indicated by yellow dashed lines). Panel **D** showed a vessel (indicated by dashed lines) with increased perivascular space containing Aβ-positive amorphous cell lysates in the lumen with some filament-like amyloid fibers starting to form (indicated by arrows). Panel **E** and **F** showed two examples of CAA with aneurysm formation (indicated by dashed lines). Panel **G** illustrated that vascular Aβ associated with abnormal swelling of vascular wall with the formation of bump-like structures (indicated by dashed lines). **H**. A capillary aneurysm (indicated by dashed lines) was observed that was directly linked to a feeding capillary (indicating by an arrow). Scale bar, 100 μm.

### Luminal and vascular buildup of Aβ associates with vascular wall swelling and aneurysm formation with the change of vascular collagen architecture

Increased perivascular space is a phenotype frequently observed in patients with Alzheimer’s diseases^31^. However, the underline cause for the “increased perivascular space” phenotype was not fully understood. We studied this phenotype further with immunohistochemistry and confocal imaging. We found that there was clear but thin ColIV immunostaining signals surrounding many vessels with “increased perivascular space”, often with concurrent perivascular Aβ deposition (Figure 10A, B, C). As we pointed out earlier, most of the small vessels with increased perivascular space had luminal Aβ staining, suggesting these vessels could be affected by blood flow blockage (Figure 9). In larger vessels with the “increased perivascular space” phenotype, we observed luminal Aβ staining, arterial dissection and increased space between split collagen layers (Figure 10D). Under some conditions, the term “increased perivascular space” might be inaccurate. At least in the cases we shown, what could be really happening is the swelling of vascular wall and increased space between split collagen layers. The phenotype with fluid leakage and swelling of the vessel wall was defined as fusiform aneurysm or saccular aneurysm in medical terms. The vascular wall swelling in the vessels with “increased perivascular space” phenotype resembles fusiform aneurysm topologically. Examples of Aβ-associated saccular aneurysm formation were shown in Figure 10D and E. We also observed that the rupture of aneurysm and the release of Aβ, which was likely an explanation for perivascular plaque formation (Figure 10F).

**Figure 10.**
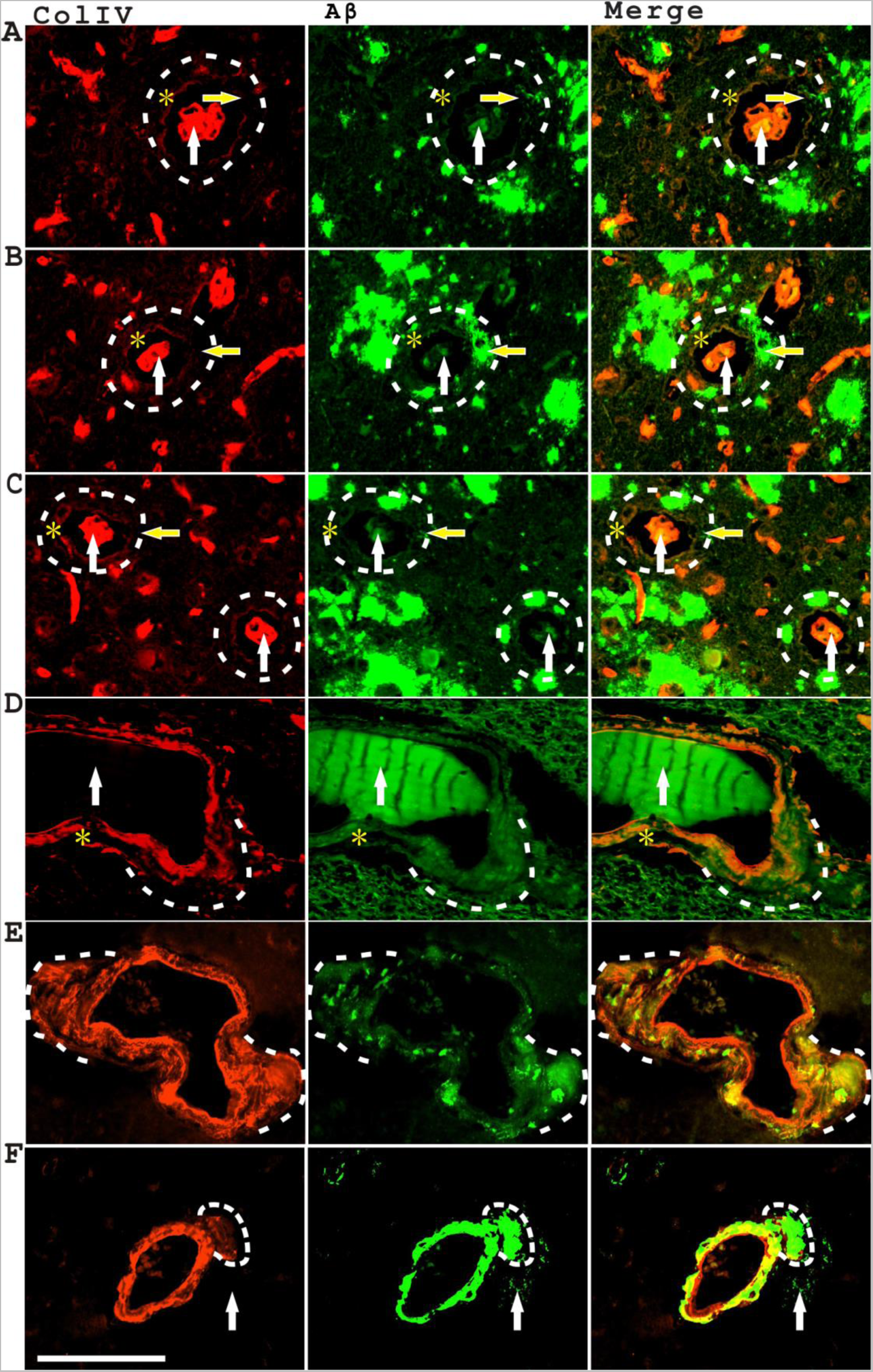
Luminal and vascular buildup of Aβ associates with vascular wall swelling and aneurysm formation. The top three panels showed 4 blood vessels (circled with dashed lines) with “increased perivascular space” (**A-C**). All these vessels showed luminal Aβ staining (indicated with white arrows) and ColIV staining in the perivascular space (indicated by asterisks) and also perivascular Aβ deposition (indicated with yellow arrows). **D**. The fourth panel showed a vascular dissection phenotype with the outer layer of vascular wall separated from the inner layer of vascular wall labeled with ColIV staining with concurrent aneurysm development (indicated by dashed lines) and luminal Aβ deposition (indicated with asterisks). **E**. The fifth panel showed additional samples of aneurysm formation (indicated by dashed lines) with vascular Aβ deposition. **F**. The rupture of aneurysm (indicated by dashed lines) linked to perivascular Aβ deposition (indicated by white arrows). Scale bar, 100 μm.

### The co-expression of Cathepsin D and Aβ was detected in senile plaques, CAA and in the blood stream of AD patients

Synthesized Aβ40 or Aβ42 peptides can easily form oligomers, or fibers if providing a proper concentration and incubation condition in test tubes *in* vitro. However, *in vivo*, Aβ peptides with amyloidogenic abilities were firstly produced from the AβPP precursor through the beta site cleavage. To understand better the mechanism of senile plaque formation, we need to know what the enzymes are responsible for AβPP beta site cleavage. During a study on the relation between lysosomes and senile plaque formation, we noticed that senile plaques had unusual staining patterns of a lysosome marker, Cathepsin D. Apparently, in senile plaques, there were two distinct layers of Cathepsin D staining. One layer was a strong, granule type of Cathepsin D staining, typical of lysosome staining, which stained complementary locations related to Aβ staining in senile plaques (Figure 11A, first, third and fifth panels). Another type was a relatively weaker and diffusive Cathepsin D expression, which often stained a large portion of senile plaque area and showed overlapping staining with Aβ (Figure 11A, second, fourth and sixth panels), which we consider as a constitutive Cathepsin D component in senile plaques. This kind of constitutive Cathepsin D staining was often detected in dense-core senile plaques. In diffusive senile plaques, the constitutive Cathepsin D expression was weak, however, still detectable. The two-layer Cathepsin D expression pattern in the senile plaques was further confirmed with a different Aβ/Cathepsin D primary antibody pair, with similar results shown in Supplementary Figure 3. Since senile plaque Aβ probably comes from blood, the constitutive Cathepsin D in the senile plaques might also come from blood leakage as well. We checked the co-expression of Cathepsin D and amyloid beta in AD patient blood vessels. At the same time, we stained the red blood cells with the histological dye Rhodanine. We did observe both Cathepsin D and amyloid beta staining associated with intravascular hemolysis in micro vessels or in capillaries on AD pathological sections (Figure 11B). Additionally, the co-expression of Cathepsin D and Aβ was also detected in CAA blood vessels (Supplementary Figure 4). Thus, Cathepsin D, a well-known candidate of beta site cutting enzyme^32^, could be an important enzyme responsible for beta site cleavage during the senile plaque formation process.

**Figure 11.**
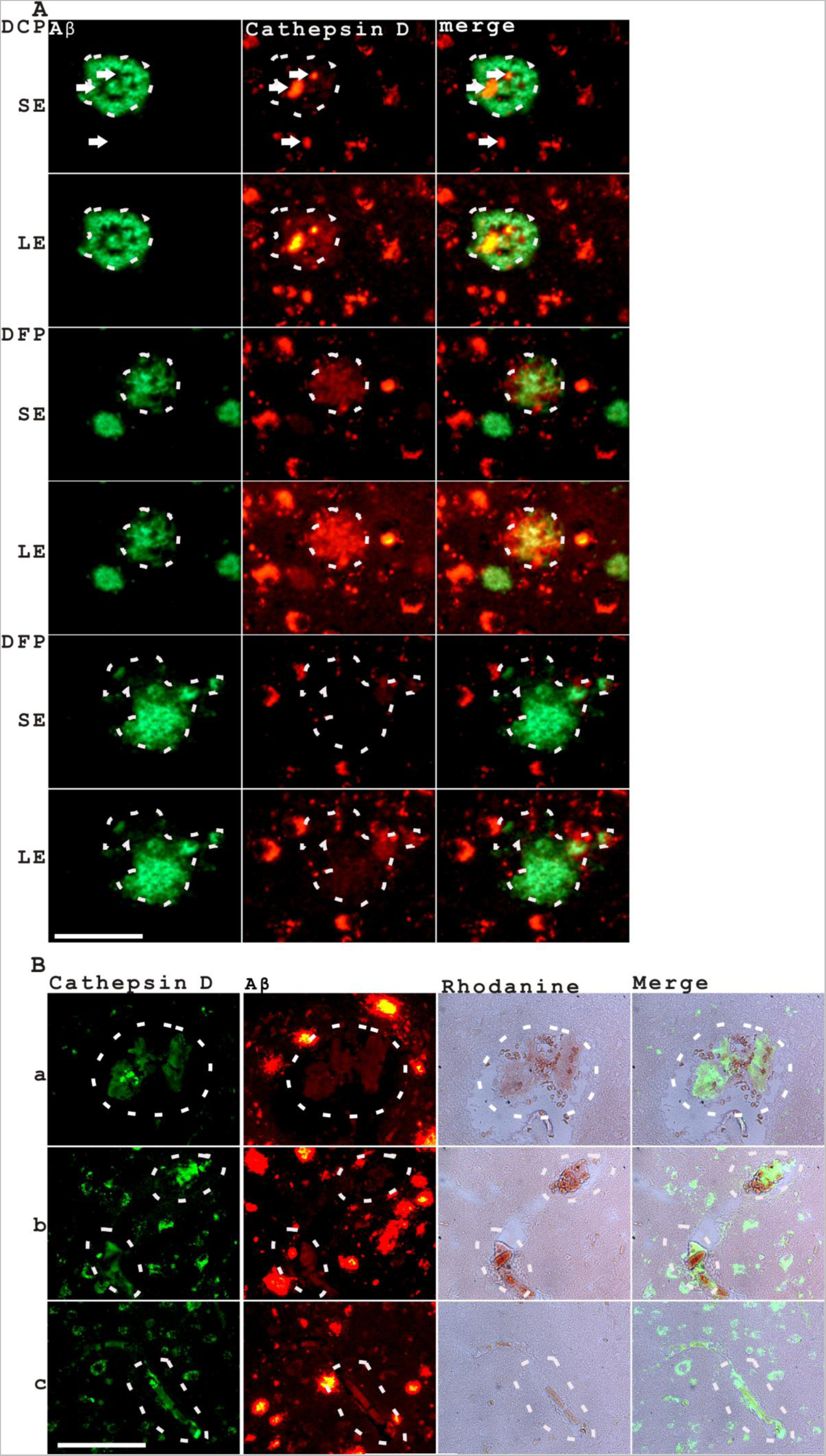
The co-expression of Cathepsin D and Aβ was detected in senile plaques and in the blood stream of AD patients. **A**. Senile plaques showed a constitutive but relatively weak Cathepsin D staining. A dense-core senile plaque (top two panels) had two distinct pattern of Cathepsin D staining. A strong and granule-like Cathepsin D staining (indicated by arrows) showed at complementary positions relative to Aβ staining with the Cathepsin D image taken at a short exposure of 0.5 second (SE). A weaker and diffusive Cathepsin D staining (indicated by dashed liens) co-distributed with Aβ staining in the senile plaques was also observed with the Cathepsin D image taken at a longer exposure of 1 second (LE) while the exposure for Aβ staining was not changed. A diffusive plaque (middle two panels) also showed a constitutive Cathepsin D staining that co-distributed with Aβ staining but not totally matching the outline of the senile plaque. A large diffusive plaque (bottom two panels) showed less detectable constitutive Cathepsin D staining. Dashed lines indicated the regions of senile plaques. The arrows indicated strong and granule-like Cathepsin D staining. Scale bar, 50 μm. **B**. Intravascular hemolysis was associated with intravascular Cathepsin D and Aβ expression in AD brain tissue blood vessels of different sizes, from medium to small vessels to capillaries (**a-c**) (indicated by dashed lines). Scale bar, 100 μm.

### Microaneurysms serve as the incubation chambers for amyloid Aβ formation

It had long been hypothesized that senile plaques are induced by cerebral microhemorrhage, however, it is not clear what kind of microhemorrhage leads to senile plaque formation. We noticed an acute RBC extravagation did not lead to typical senile plaque formation instantly(Supplementary Figure 5). It is possible that senile plaque formation is driven by a chronic microhemorrhage process. We previously observed primitive senile plaque development was associating with hemoglobin, ApoE and Cathepsin D expression, suggesting that the preplaque formation was associated with hemolysis^24^. In this study, by using phase-contrast or DIC objective lenses, we further found that the primitive plaque formation was actually happening within membrane-enclosed sac-like structures. These sac-like structures had an average diameter of 43.96 (±19.79) μm (N=39), which is not statistically different from the size of an average senile plaque with the diameter of 38.67 (±15.94) μm (N=125, p=0.13). Since these saccular structures contained blood and vascular markers such as hemoglobin and ACTA2, they are most probable microaneurysm structures. We were astonished to find that some of our microaneurysm images (For example, an microaneurysm image in Figure 9H) were strikingly similar to early descriptions on Charcot-Bouchard aneurysm by Charles Jacques Bouchard, who pioneered the works that linked Charcot-Bouchard aneurysm to cerebral microhemorrhage^24,25^. Apparently, Charcot-Bouchard aneurysm is mostly linked to a chronic form of cerebral microhemorrhage.

We went through our pathological images to look for more evidence showing the link between microaneurysm and senile plaque formation. We could observe several to hundreds of saccular microaneurysms in different AD pathological sections. These saccular microaneurysms contained Aβ, blood or plasma markers such as MetaBlue, HBA, ApoE, vascular markers such as ACTA2, and the important beta-site enzyme Cathepsin D (Figure 12). Some of these microaneurysms contain diffusive amorphous Aβ resembling primitive diffusive plaques (Figure 12B). Some other samples contained Aβ staining with a denser core resembling primitive dense-core plaques (Figure 12C, D). In some plaques with typical mature dense-core senile plaque morphology, the saccular membrane structures still persisted (Supplementary Figure 6). A relatively intact core of a saccular microaneurysm or a blood vessel might be the structure base for the cores of dense-core plaques while the central membrane structures degenerated in diffusive plaques. In many of these microaneurysm membrane structures, dark materials under phase contrast objective lens were also observed (Figure 7A, D, E, F), likely by-products of vascular/hemolytic degeneration.

**Figure 12.**
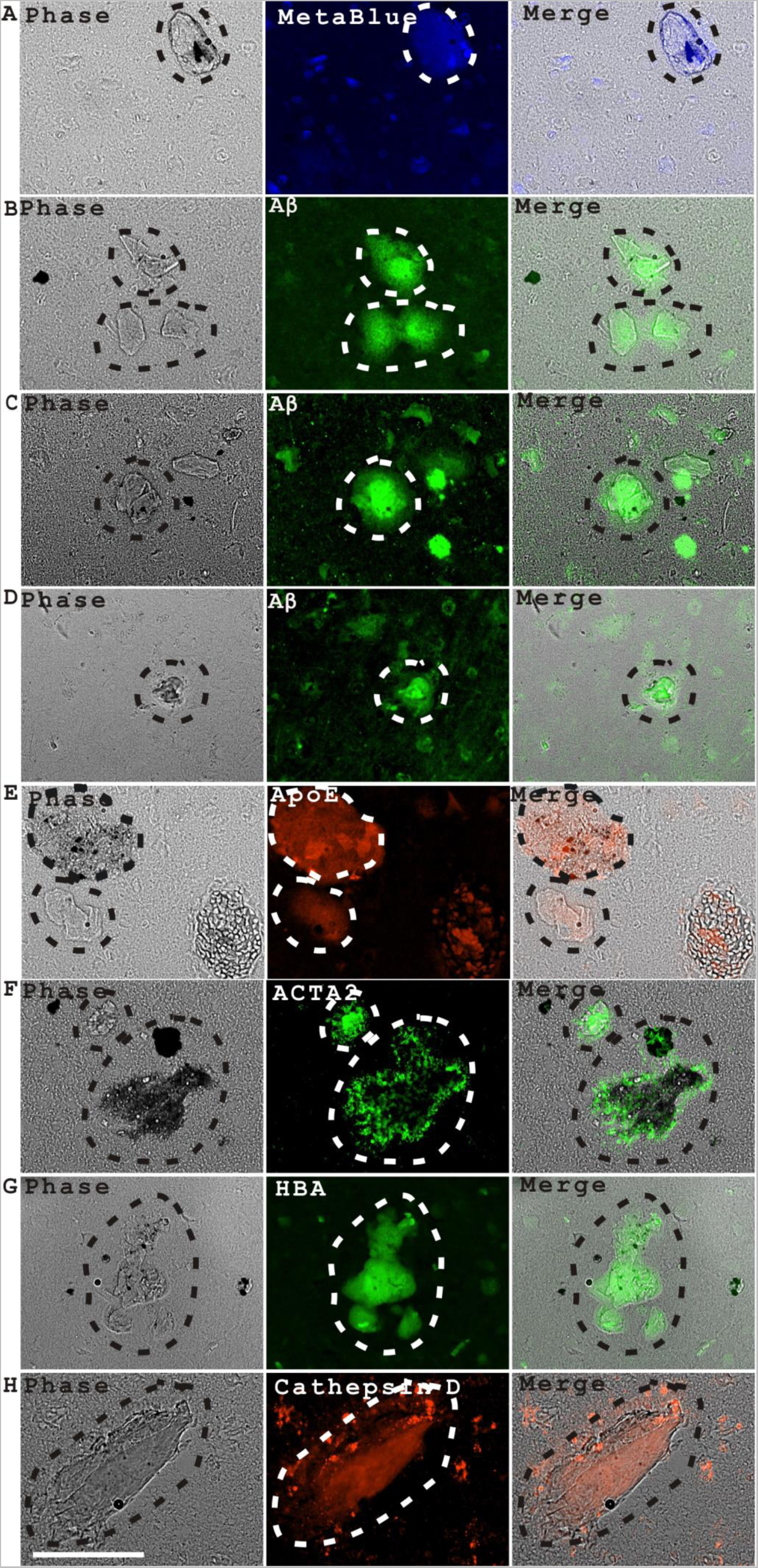
Saccular aneurysms serve as the incubation chambers for Aβ polymerization and preplaque formation. Saccular aneurysms were marked with the blood, vascular and lysosomal markers such as MetaBlue, Aβ, ApoE, ACTA2, HBA and Cathepsin D. **A**, MetaBlue; **B-D**, Aβ; **E**, ApoE; **F**, ACTA2; **G**, HBA; **H**, Cathepsin D. Scale bar, 100 μm.

In some AD sections, we observed even more severe type of vascular degeneration than microaneurysm degeneration as shown in Supplementary Figure 7. In some cases, long stretches of blood vessels degenerated with diffusive HBA and Aβ staining (Supplementary Figure 7A). In other cases, the total degeneration of large vessels was observed accompanying with perivascular HBA and Aβ staining (Supplementary Figure 7B). Occasionally, the simultaneous degenerations of multiple tangled blood vessels in clusters were also observed (Supplementary Figure 7C, D). Dark materials surrounding these degenerated blood vessels were also observed in this figure (Supplementary Figure 7A, B and C).

### Aβ forms protease-sensitive heterooligomers rapidly with hemoglobin while Aβ self-oligomers are protease-resistant

Since the senile plaque formation process might start from hemolysis, we thought the initial Aβ aggregation might also have something to do with hemolysis. We hypothesized that HBA physically interacts with Aβ during hemolytic events. The potential interaction between hemoglobin and Aβ has been studied in the past in a few published studies^33,34^, but the investigations were still not complete and comprehensive. We studied the Aβ/Hb interaction with an aggreagation assay in the test tubes. First, we checked the self-oligomerization of Aβ peptides (Figure 13). Both Aβ40 and Aβ42 formed small aggregates immediately soon after dissolving the peptides in DMSO and diluted into PBS solution at 5 μM concerntration. Aβ40 formed aggregates with increasing size upon incubation up to 3 days, while the sizes of Aβ42 self-oligomer did not change significantly upon prolonged incubation (Figure 13A). Both Aβ40 and Aβ42 aggregates were resistant to a 15-minute Proteinase K (PK) treatment, suggesting the self aggregates were protease-resistant (Figure 13B).

**Figure 13.**
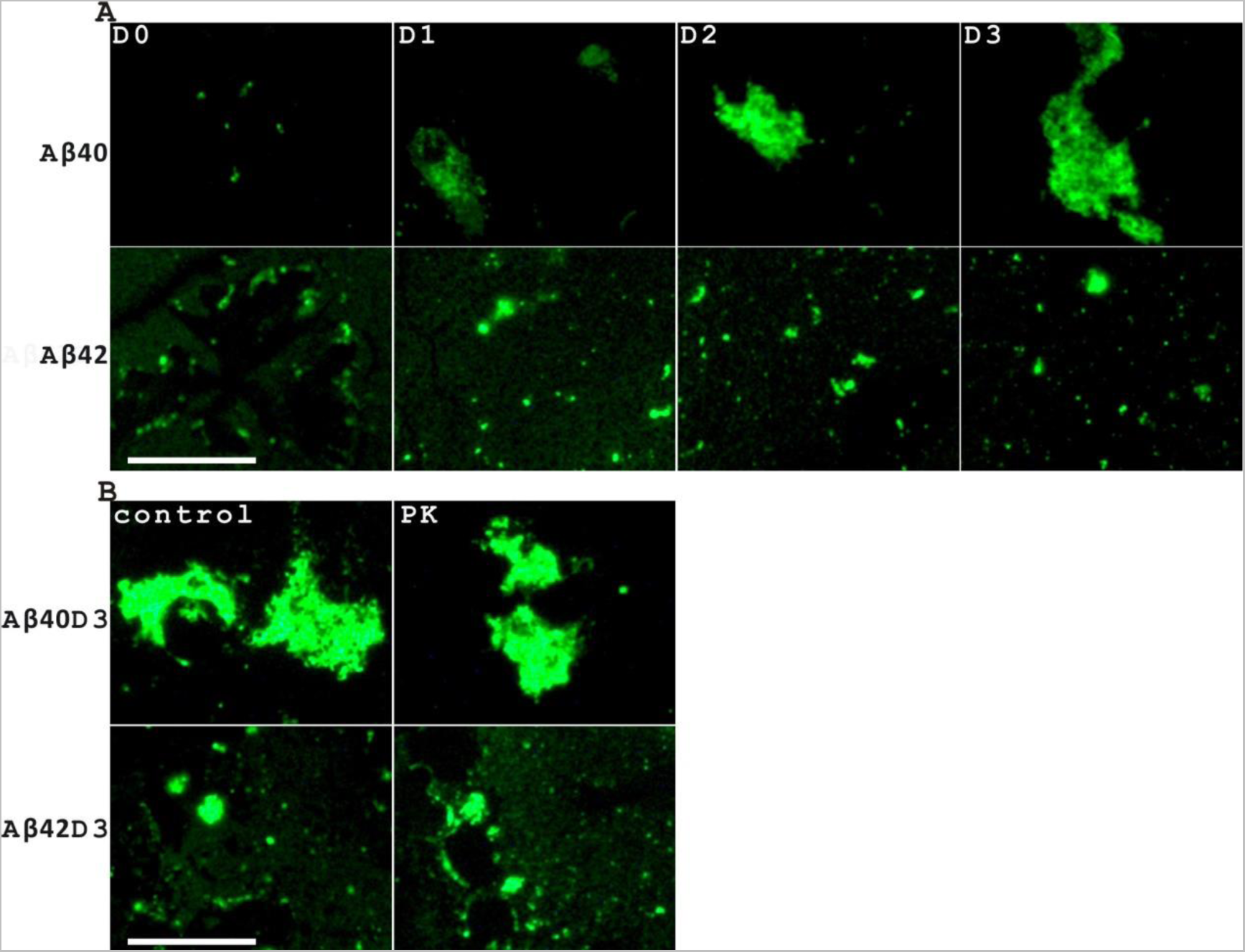
Aβ40 and Aβ42 form protease-resistant self-oligomers *in vitro*. **A.** Aβ40 and Aβ42 formed self-oligomers when incubating in PBS solution for up to 3 days (Day 0 to Day 3) followed with a drying up procedure by droplet evaporation. D0 stands for Day 0. **B.** Both Aβ40 and Aβ42 self-oligomers were resistant to Proteinase K (PK) treatment. Scale bar, 50 μm.

We then incubated hemoglobin and Aβ40 at a molar ratio of 1:4 (1.25 μM hemoglobin and 5 μM Aβ40), and checked the aggregation product out of this mixture. We observed that hemoglobin formed complexes with Aβ40 very quickly. Even in Day 0 samples, hemoglobin formed large leaf-like or ring-shape complexes with Aβ40 (Figure 14A, top two panels). Some dot-like structures dervied from Aβ40 self-oligomerization were also present in Day 0 samples (Figure 14A, third panel). Large leaf-like structures and ring-shape structures were continually present in Day 1, Day 2, Day 3 samples (Figure 14A, bottom four panels). In AD tissue samples, we did not observe large-leaf like structures with Aβ staining. However, we did observe some ring-shape structures with overlapping staining of Aβ and hemoglobin in blood vessels with intravascular hemolysis (Figure 14B). which suggested that the ring-shape Aβ/Hb complexes could form in the blood vessels of AD tissues *in vivo*. We did more studies on the Aβ/Hb interaction on red blood cells on pathological sections as shown in Figure 17.

**Figure 14.**
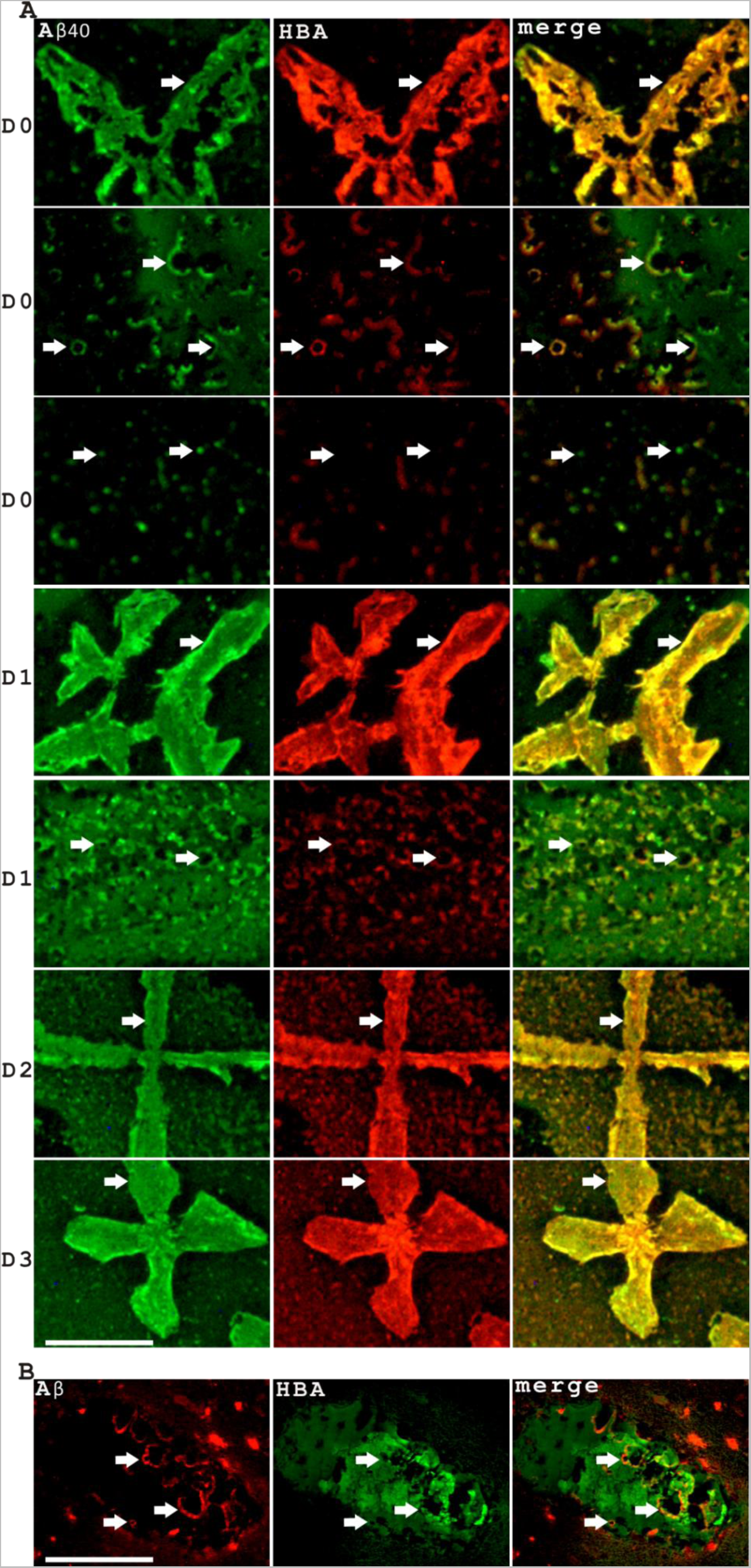
Aβ40 forms heterooligomers rapidly with hemoglobin. **A.** Aβ40 forms heterooligomers rapidly with hemoglobin when co-incubating in PBS solution for up to 3 days followed with sample drying by droplet evaporation. Co-immunostaining of Aβ and hemoglobin was shown for Day 0 (top 3 panels), Day 1 (fourth and fifth panels), Day 2 (the sixth panel) and Day 3 (the seventh panel) samples. Arrows indicated different structures, such as leaf-shape, ring-shape structures, that co-stained with both Aβ and HBA (alpha hemoglobin) antibodies. The dot shape structure likely represented Aβ40 self-oligomers. Some ring shape structures were indicated with arrows in the second and fifth panel. Scale bar, 50 μm **B.** Aβ forms ring-shape structures interacting with hemoglobin in the lumens of AD patient blood vessels similar to the ring-shape structures formed in Aβ-and-hemoglobin mixture samples *in vitro*. Scale bar, 100 μm.

In Aβ42 and Hb mixed Day 0 samples, we observed leaf-shape, ring-shape, and line-shape structures with both Aβ42 and Hb staining (Figure 15). In Day 0 samples, we also see the existence of Aβ42 self-oligomers, which was less frequently detected in Day 1, Day 2 and Day 3 samples (Figure 15A), indicating that Aβ42/Hb interactions might compete out Aβ42 self-oligomerization. Previous studies suggested that Hemin inhibits Aβ aggregation^35,36^ and hemoglobin might interact with Aβ through the Hemin motif^34^, suggesting that a hemoglobin inhibition of Aβ self-oligomerization is likely. Large leaf-shape or smaller ring-shape structures were continually observed in Day 1, Day 2 and Day 3 samples (Figure 15, bottom three panels). Unlike the protease-resistant Aβ40 and Aβ42 self-oligomers, Aβ40/Hb and Aβ42/Hb hetero-complexes were both sensitive to PK treatment (Figure 15B).

**Figure 15.**
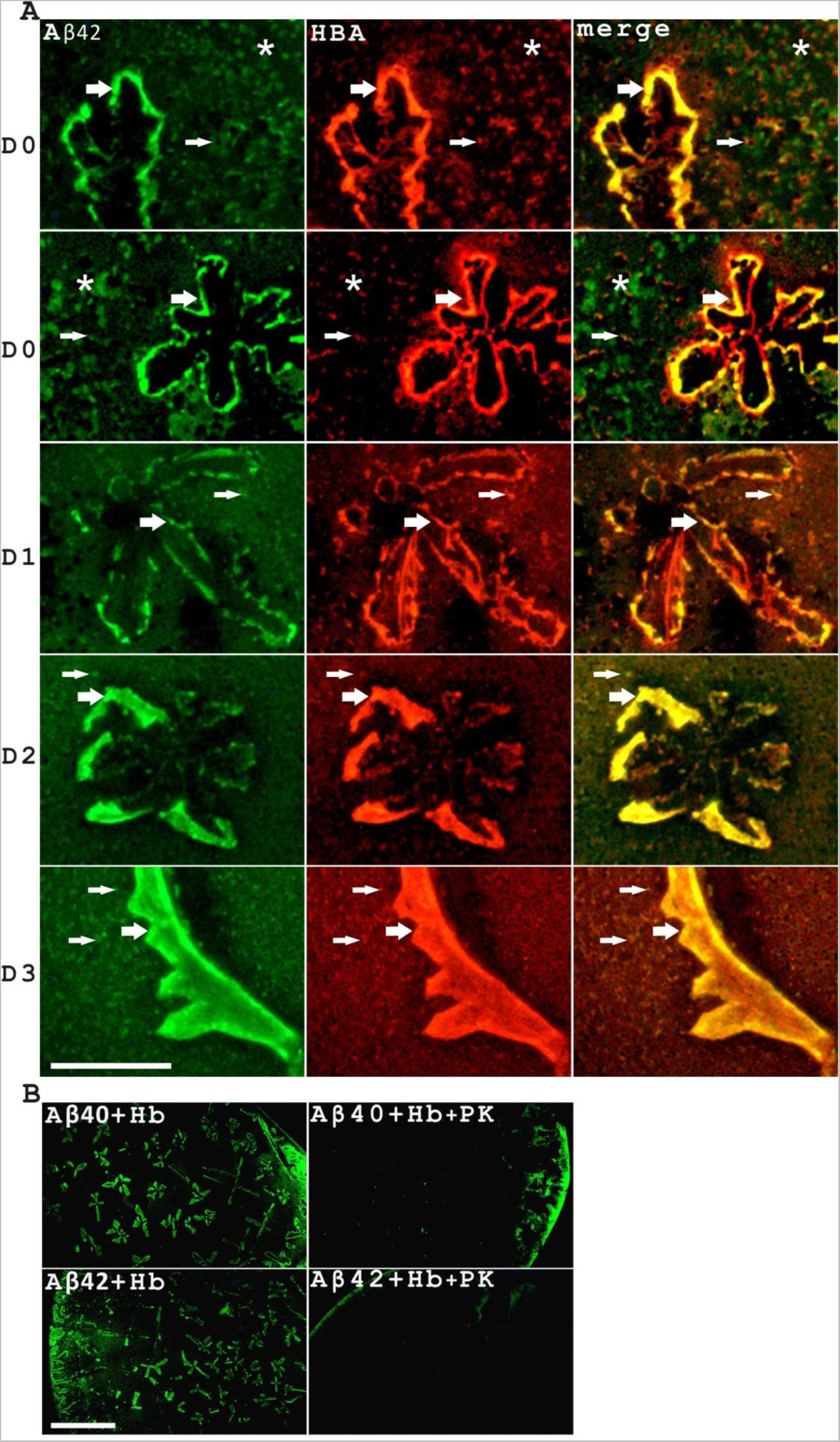
Aβ42 forms protease-sensitive hetero-oligomers rapidly with hemoglobin when co-incubating in PBS solution for up to 3 days followed with sample drying by droplet evaporation. **A.** Aβ42 forms heterooligomers rapidly with hemoglobin in PBS solution. Co-immunostaining of Aβ and hemoglobin was shown for Day 0 (top two panels), Day 1 (third panel), Day 2 (fourth panel) and Day 3 (fifth panel) samples. Large arrows indicated relatively large structures while small arrows indicated smaller structures. Leaf-shape, ring-shape or dot-shape structures were observed. Some self-oligomers also existed in Day 0 samples, indicated with asterisks. Scale bar, 50 μm. **B.** Both Aβ40/Hb and Aβ42/Hb complexes were protease-sensitive with the complexes illustrated by Aβ antibody staining. PK treatment drastically decreased the signals of resulting complexes. Scale bar, 200 μm.

### There is a specific Aβ interaction with hemoglobin at single cell levels in red blood cells

The *in vitro* hemoglobin/Aβ binding assay indicated that Aβ might interact with hemoglobin directly, which intrigued us to look for more evidence of *in vivo* Aβ-hemoglobin interactions. We did more Aβ and HBA co-immunostainings and focused on their co-expression in red blood cells. The expression of Aβ in RBC is not constitutive since a lot of RBCs did not have Aβ expression (Figure 16A). We found at least two patterns of Aβ staining on red blood cells with one dominant type of staining co-localizing with hemoglobin while another type of staining showing no specific interaction with hemoglobin. Most often, Aβ showed as dots, stripes or diffusive patterns on red blood cells and co-localizing with hemoglobin staining (Figure 16B, C). RBCs with the stripe pattern look like fragmented RBCs, resembling helmet cells that are a type of red blood cells under stress. Additionally, we also observed Aβ/Hb staining as curved lines on red blood cells with abnormal morphology resembling schistocytes, another type of stressed red blood cells (Supplementary Figure 8). It is not clear what give rises to another less frequent pattern of Aβ-RBC interaction showing as ring-shape Aβ staining encircling red blood cells but in a way not specific to hemoglobin co-localization (Figure 16D, E). It is possible that multiple species of Aβ related peptides or peptide complexes might give rise to different patterns of RBC interactions. The results also showed that Aβ staining was enhanced in hemolysis, further emphasizing a role of the Aβ-hemoglobin interaction in RBC stress (Figure 16F, G and H). In some vessels with intravascular hemolysis, an Aβ staining pattern independent of hemoglobin interaction was also noticed, which could be due to the formation of Aβ self-oligomers instead of Hb/Aβ heterocomplexes.

**Figure 16.**
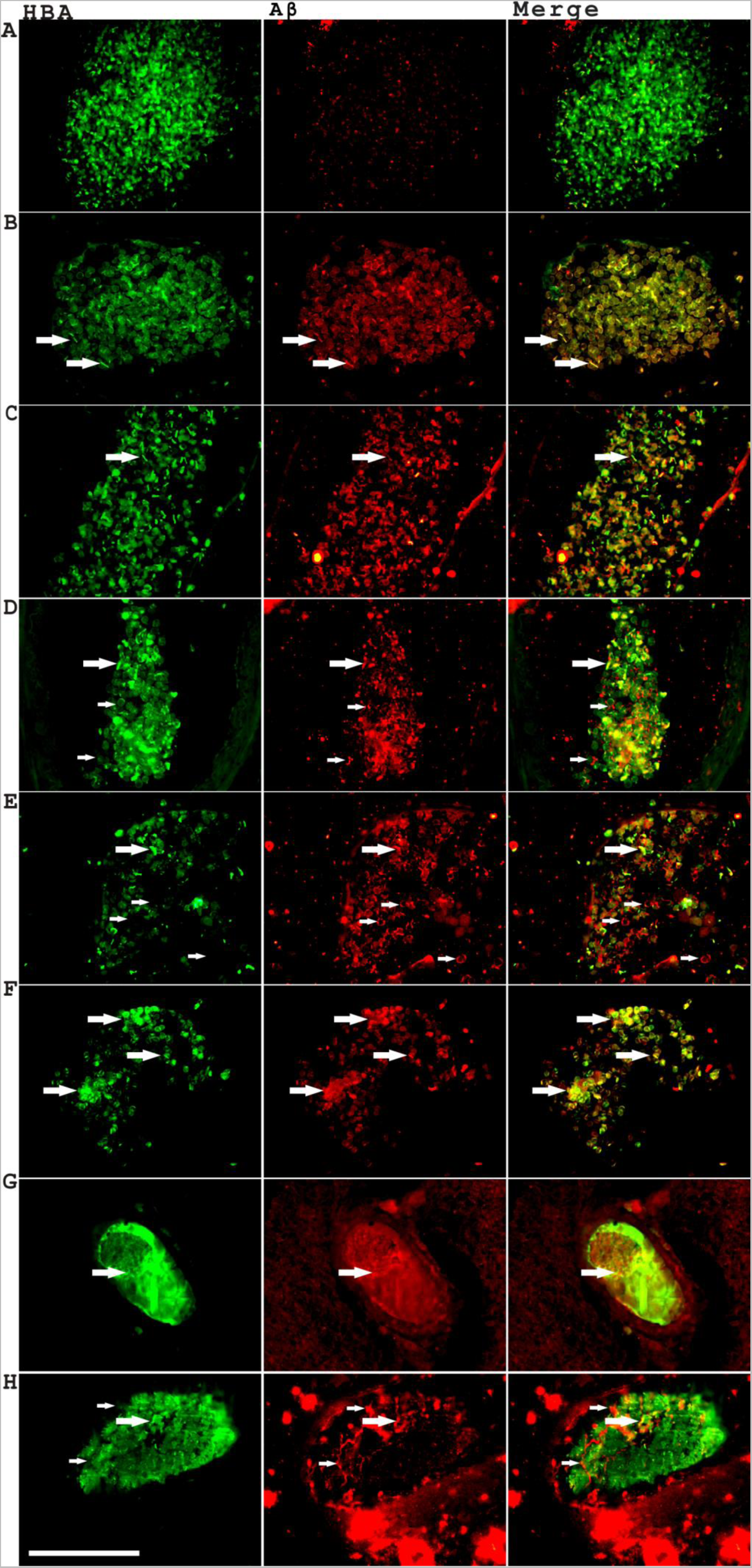
Aβ interacts with hemoglobin at single cell levels in red blood cells. **A.** The top panel showed that many RBCs are lack of Aβ staining, which means that the association between Aβ and RBC is not constitutive. (**B, C**): The second and third panel showed that Aβ can associate with HBA in dots, stripes or diffusive forms (indicated with large arrows). (**D, E**): The fourth and fifth panel showed that Aβ sometimes associates with RBC in a non HBA-specific manner (indicated with small arrows). (**F-H**): The sixth, seventh and eighth panels showed that significant Aβ staining was associated with intravascular hemolysis with hemolytic HBA staining patterns. In panel H, a type of strong Aβ staining not co-localizing with hemoglobin was also observed (indicated with small arrows). Scale bar, 100 μm

**Figure 17.**
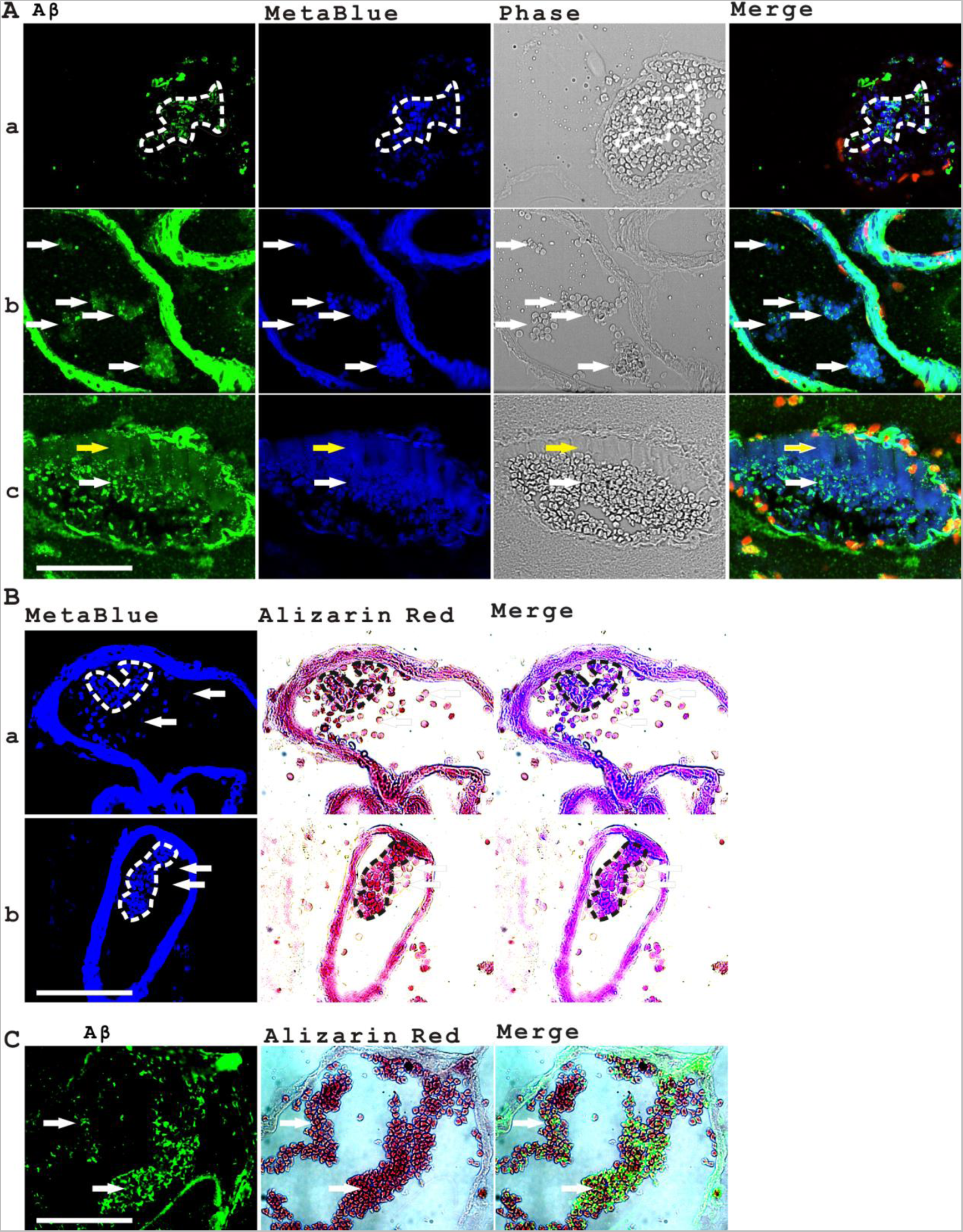
The interaction between MetaBlue and Aβ associates with RBC coagulation and calcium signaling. **A**. The top panel showed that Aβ staining associated with MetaBlue staining in RBCs at the center of aggregating red blood cells (**a**). The middle panel showed the Aβ-MetaBlue association in disseminated intravascular coagulation of red blood cells (**b**). The third panel showed that Aβ was associated with intravascular hemolysis and aggregating RBCs (**c**). The dashed lines and the white arrows indicated coagulating RBCs. The yellow arrow indicated hemolysis. **B**. MetaBlue autofluorescence was specifically enhanced in RBCs with higher calcium staining by Alizarin Red, which is also a marker of RBC coagulation. The dashed lines indicated coagulating RBCs (**a, b**). The arrows indicated that the red blood cells with weaker MetaBlue fluorescence also had weaker calcium level shown by Alizarin Red staining. **C**. A significant expression of Aβ was detected in aggregating RBCs marked with Alizarin Red. Arrows indicated coagulating RBCs with Aβ staining. Scale bar, 100 μm

The specific interaction between Aβ and hemoglobin both *in vivo* and *in vitro* raises another important question: what is the real physiological significance of this interaction? With confocal microscopy, we found that both Aβ and MetaBlue were associating specifically with aggregating red blood cells in the center of large red blood cell clusters (Figure 17A, top panel) or in disseminated intravascular coagulations (Figure 17A, middle panel). Significant Aβ and MetaBlue staining was also linked to intravascular hemolysis (Figure17A, bottom panel) as previously already shown. Since both vascular aging and red blood cell coagulation has been linked with calcium signaling, we stained the pathological sections with the calcium specific dye Alizarin Red. Alizarin Red staining showed clearly that CAA vessels were calcified (Figure 7H). Alizarin red also labels RBCs very well suggesting that RBCs are critical sources of calcium *in vivo*. In aggregating RBCs, Alizarin Red staining was intensified with simultaneous enhancement of MetaBlue autofluorescence (Figure 17B). Thus, MetaBlue fluorescence is linked to calcium-regulated RBC coagulation. Alizarin Red staining also clearly showed that Aβ staining was enhanced in aggregating red blood cells (Figure 17C).

## Discussion

Senile plaque Aβ has been hypothesized to be derived from neural or vascular/blood source^7,12,15,37^. This study indicated the primary direct source of Aβ is not neural cell with the evidence showing a lack of substantial overlapping of senile plaque Aβ with neural markers. Additionally, Aβ expression in neural cells appears to be too low to account for the high density Aβ expression in senile plaques. Moreover, compact plaques and the cores of dense core plaques did not have nuclei or TUNEL-labeled nuclei, not supporting that senile Aβ comes from dying neural cells. Instead, senile plaque Aβ most likely comes from hemolysis-associated Aβ leakage out of damaged blood vessels. The evidence is the following: Firstly, RBCs in AD patients were clearly stained with Aβ antibody, especially in coagulating or hemolytic RBCs with clear co-localization with hemoglobin in the patterns of dots, stripes or diffusive patterns. Secondly, strongly-stained Aβ aggregates were present in the hemolytic regions. Thirdly, Aβ co-distributed with multiple blood or plasma markers such as HBA, Hemin, ApoE and MetaBlue autofluorescence at multiple stages of senile plaque development such as CAA, dense-core plaques and diffusive plaques. Fourthly, the cores of dense-core plaques were wrapped with vascular or astral glial components outside and loaded with blood contents inside, indicating that the dense cores are highly organized vascular structures. Finally, microaneurysms were identified as major sites of amyloid formation besides CAA and regions of intravascular hemolysis. Microaneurysm formation and rupture was associated with Aβ polymerization and release into brain parenchyma and directly contributes to the morphogenesis of dense-core or diffusive senile plaques in AD tissues. Microaneurysm-associated brain microhemorrhage has been studied since 1865 when Bouchard and his supervisor **J**ean-Martin Charcot pioneered their studies on Charcot-Bouchard aneurysm. It came as a complete surprise that this type of aneurysm and microhemorrhage links to AD senile plaque development, so that we put “Charcot-Bouchard aneurysm” in the article title to show our respect for their ground-breaking work. Interestingly, Bouchard originally described the aneurysms as “small spots of a violaceous, blackish, or ochre color, globular in form, and varying in size from that of a millet seed to that of a pin’s head.”^30,38^. Although the microaneurysms we measured here probably were even smaller, we also detected a type of “blackish” materials in the microaneurysms. This type of blackish materials co-distributed with Aβ, HBA, Hemin, MetaBlue and Alizarin Red staining in the microaneurysms and could also be detected in some senile plaques, adding another evidence that senile plaques arise from microaneurysm degeneration (Figure 7 and Supplementary Figure 7, 9, 10). The blackish materials could be byproducts of vascular/blood degeneration with their identities worthy of further investigation.

Although mounting researches today focused on the biological effect of Aβ in neurons or glial cells in AD, however, AβPP was found to have coagulation modulation function early on, which potently inhibits platelet coagulation factor XIa, under an alternative gene name as Nexin2^31^. Another related research indicated that Aβ release is linked to thrombosis^39^. In the present study, we found that Aβ associated with multiple blood and vascular phenotypes such as RBC stress, coagulation, hemolysis, CAA, increased perivascular space, microaneurysm, and atherosclerosis (vascular plaques, blood flow blockade and vascular calcification are major characters of atherosclerosis.), highlighting an important role of Aβ in the disrupted blood homeostasis and circulation, which are fundamental defects in AD patients. This study provided both histological and biochemical evidence that Aβ might interact with hemoglobin physically. Aβ staining is enhanced in coagulating or hemolytic RBCs, which suggested an important role of Aβ in red blood cell stress. It is possible that RBC damage and hemoglobin denaturation in coagulating or hemolytic RBCs attracts Aβ to hemoglobin. The interaction between Aβ and Hb could be very important as previous research suggested that Aβ binding reduces Hemin toxicity^34^. Our experiment showed that Aβ/Hb complex was sensitive to protease digestion but Aβ self-oligomers were protease resistant, which explains why Aβ stands out as the everlasting and more stable component of senile plaques.

It is not a surprise observation that Aβ-ApoE interaction induces vascular amyloid plaques that restrict blood flow in capillaries and small vessels. There is a long-hold “hypoperfusion hypothesis” states that inadequate blood supply is a main culprit behind Alzheimer’s Disease^40,41^. Our observations clearly supported this hypothesis. Recently, there were some excellent researches showing the vascular amyloid production and an increased risk of newly-onset Alzheimer’s disease in Covid-19 patients^42–44^. In addition, the development of brain amyloid deposits in young patients with Covid-19 infections was also reported^45^. These new observations aligned well with our study that blood flow blockade by vascular amyloid plaques might be linked to senile plaque formation in the brain and AD pathogenesis. These studies suggested that vascular or blood-related infections might put patients at high risk of AD development. Together, these data emphasized that Aβ and ApoE could be both important modulators of blood homeostasis.

Two critical aspects of senile plaque formation are the source of Aβ peptides and the source of beta-site cutting enzymes. This study indicated that hemolysis, CAA and microaneurysm could be critical sites that bring together Aβ-related peptides and Cathepsin D, a potent beta-site cutting enzyme. Platelet could be an important source of Aβ-related peptides in blood coagulation and hemolysis since it is well-known that platelets secrete large amount of AΒPP in alpha-granules upon activation. Previous studies already found that Cathepsin D co-localized with senile plaques, while the source of Cathepsin D had been claimed as neuronal^27,46–48^. However, previous publications also showed that Cathepsin D could be extracellularly released as a part of lysosome exocytosis by various cell types, including platelets^49^, mast cells^50^, neutrophils^51^, macrophages ^24^or cytotoxic lymphocytes^52^ under stress conditions, which could be a way to supply Cathepsin D extracellularly in the amyloidogenic process. Along with other factors that might affect blood vessel permeability such as histamine^53^, the Cathepsin D-and-Aβ enriched hemolytic mixture could disrupt the vasculature, which induces further leakage, extracellular matrix digestion, the formation of CAA and microaneurysm. Given the long and complicate senile plaque formation process and multiple cell types involved, the participation of other lysosomal or non-lysosomal proteases such as BACE1^54,55^, Cathepsin B^56,57^, Carboxypeptidase B^58^ or MMP2^59^ should not be excluded at the present. It is quite likely that the seemingly extreme simple and ordered structures of senile plaques actually come from a complicated “chaotic” process that cannot be simply described with linearized enzyme-substrate reaction and polymerization process.

Previous studies in AD patients and in mouse models of AD have already found that dense-core plaques are centered on vessel walls^13,14^. These studies suggested that perturbed vascular transport and/or perivascular enrichment of neuronal Aβ leads to the formation of vasocentric dense plaques. Our results showed that many dense-core plaques are intermingled with degenerated vessel fragments by ColIV and LRP1 staining, providing further evidence that vascular degeneration is intrinsically linked with senile plaque formation. However, we think that senile plaque Aβ is primarily derived from the blood source although the additional contribution of Aβ from vascular cells or neural cells is possible. A series of studies by Dr. Cullen’s group using thick brain tissue sections showed a clear anatomic link between microhemorrhage with senile plaque formation and also Tau phosphorylation^12,17,18^.

Yet, in these studies, the exact source of Aβ and how exactly microhemorrhage leads to senile plaque development was still not clear. With Rhodanine and Alizarin Red staining instead of Perls staining and also Hb immunohistochemistry, our study provided a direct link between blood Aβ expression and RBC coagulation or hemolysis, suggesting that stressed or damaged RBCs were enriched for Aβ. In addition, the data suggested that CAA formation funnels the blood or plasma Aβ into the vascular wall, promoting the formation of microaneurysms, which are the major sites of preplaque formation. In summary, our data suggested that the senile plaque formation is primarily induced by the leakage of Aβ-associated hemolysates with CAA and microaneurysm-associated vascular degeneration as intermediate steps. This new theory of vascular/blood origin of senile plaque Aβ explained well on the new data, such as the presence of hemoglobin, Hemin and MetaBlue fluorescence in the senile plaques, the nucleus-null dense cores, microaneurysmal amyloid formation, and the ring-shape or asymmetrical Aβ staining in the dense cores and the dense-core saccular membrane structures etc.

The findings that senile plaques are associated with specific blue autofluorescence are intriguing. There have been a number of studies related to senile plaque blue autofluorescence materials (temporarily named as MetaBlue in this article) in the past^22,23,60^. An early study reported that the blue autofluorescence was associated with senile plaques but not CAA^22^. Two more recent studies found that the blue autofluorescence was associated with both senile plaques and CAA^23,60^. Our study additionally found that MetaBlue was not only associated with CAA and senile plaques but also directly associated with red blood cells, adding another evidence that senile plaques come from blood/vascular origin. It should be emphasized that MetaBlue fluorescence is an intrinsic blue fluorescence, not dependent on blue fluorescent nuclei dye staining such as Hoechst staining, thus previous publications using blue fluorescent nuclei dye in the AD research probably should have their nuclei staining data re-examined. Our study provided a link between MetaBlue and RBC coagulation and also the calcium level in RBC for the first time, suggesting MetaBlue probably has an important physiological function in RBC physiology. The exact identity of MetaBlue is unknown. MetaBlue could be an oxidative-stress related metabolite since previous literature indicated that the blue autofluorescence of amyloid peptides could be associated with tyrosine oxidation^61^. To isolate MetaBlue materials and to elucidate the true identity of MetaBlue will likely leads to deeper understanding of AD pathogenesis.

## Data and material availability

All data needed to evaluate the conclusions in this paper are present either in the main text or in the supplementary materials.

## Supplementary Materials

Supplementary Figures 1-10

## Acknowledgements

We want to thank the great help from Dr. Ma Chao and Dr. Qiu Wenying for providing AD tissue sections from National Human Brain Bank for Development and Function, Chinese Academy of Medical Sciences and Peking Union Medical College, Beijing, China. Additionally, we want to thank Dr. Xiangli Zhang of Shandong University and Dr. Chunlei Zhang of Shanghai Jiao Tong University for valuable discussions. We also want to thank Shunjie Wu from Keyence Co. for allowing us to use their Keyence BZ-X800 microscope without charge. Furthermore, we want to thank Housheng Wang, Lu Xu, Wenting Xuan, Pingxin Liu, Xiaoyi Bao, and Jun Wang for excellent lab assistance.

## Author contributions

H. F. conceived the study; H. F. designed and supervised the experiments; H. F. and J. L. performed the experiments; H. F., J. L., P. D., W. J., G. G., D. C. did the data analysis; H. F. wrote the manuscript; All authors reviewed the manuscript.

---

**Supplementary Figure 1.**
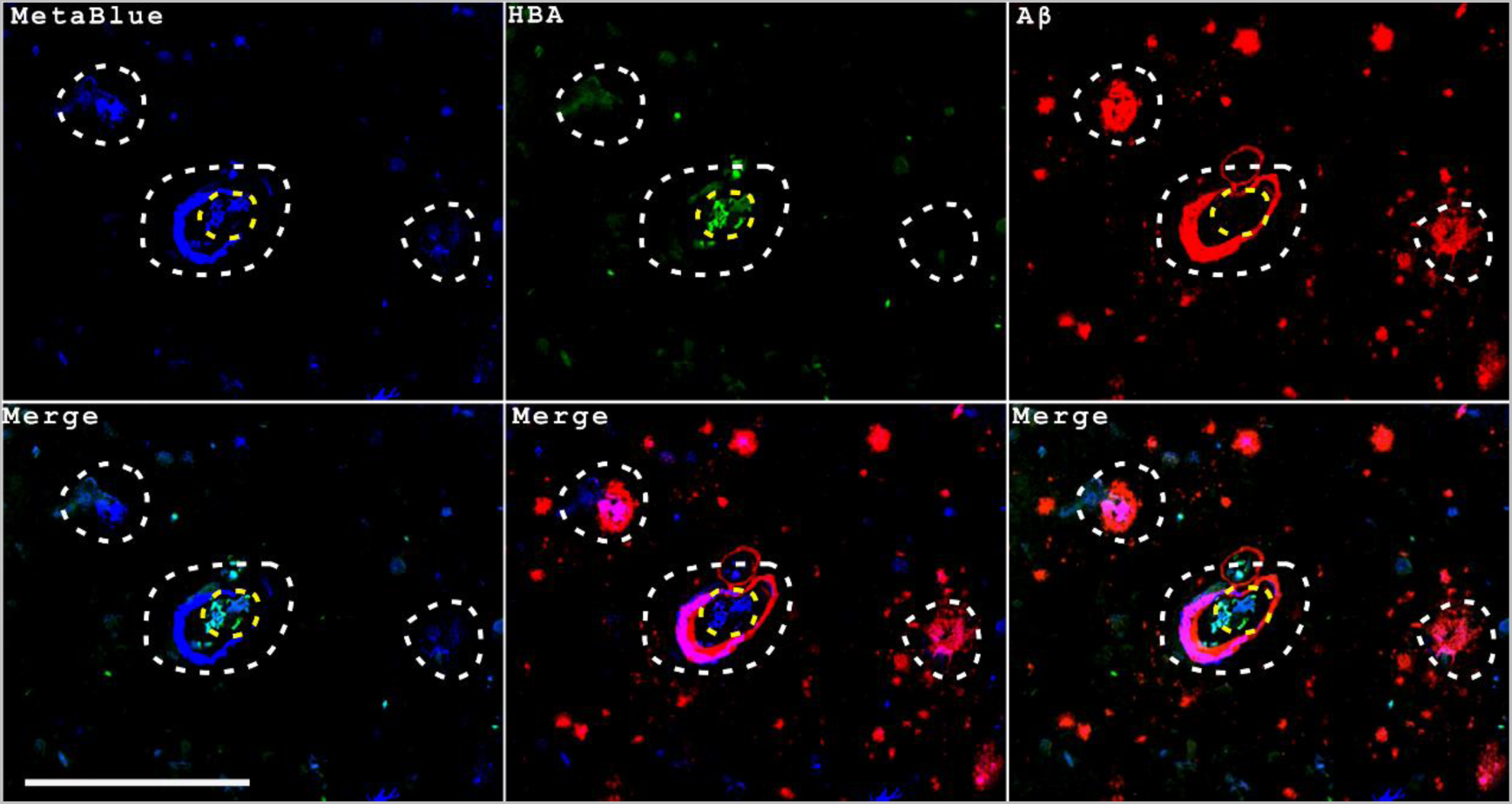
MetaBlue blue fluorescence is present in CAA, dense-core plaques, diffusive plaques and also red blood cells. The white dashed lines indicated a dense-core plaque, a CAA blood vessel and a diffusive plaque as labeled by Aβ (from left to right). The yellow dashed lines indicated red blood cells as labeled by HBA. MetaBlue fluorescence was present in all these structures with variable intensities. Scale bar, 200μm.

**Supplementary Figure 2.**
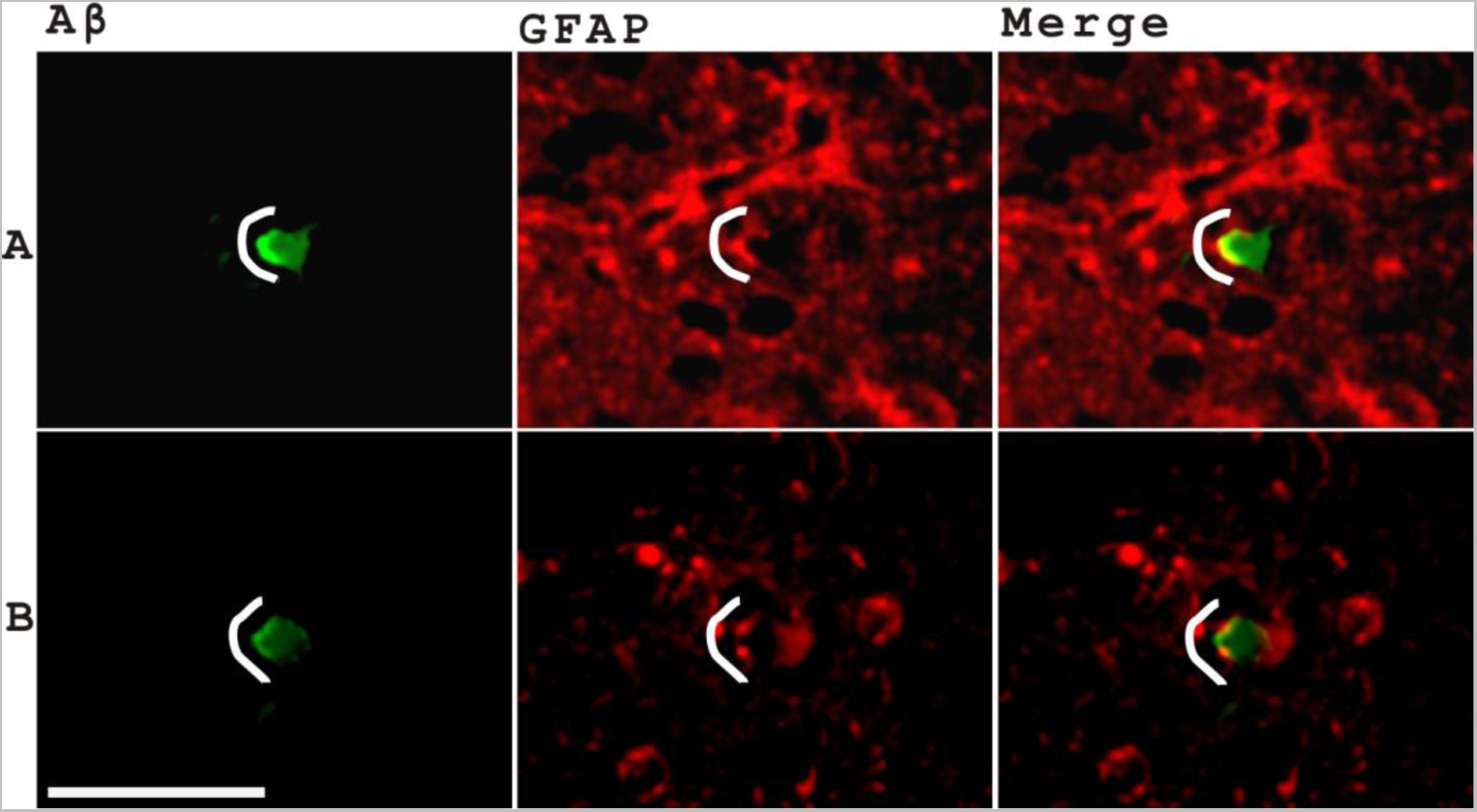
Asymmetrical distribution of Aβ and glial foot processes with the core barrier structure half broken suggested the Aβ flow direction is from inside to outside of the core. The two panels showed two dense cores with half broken glial foot barrier structures (**A, B**). The half with intact glial foot structures had stronger Aβ immunostaining than the half losing the glial foot structures. The exposure time for the green channel was shortened in order to show the fine details of asymmetrical Aβ intensity distribution of the dense cores so the rims of these plaques were less visible. The white lines indicated the halves with relatively intact glial foot structures surrounding the dense cores. Scale bar, 50μm.

**Supplementary Figure 3.**
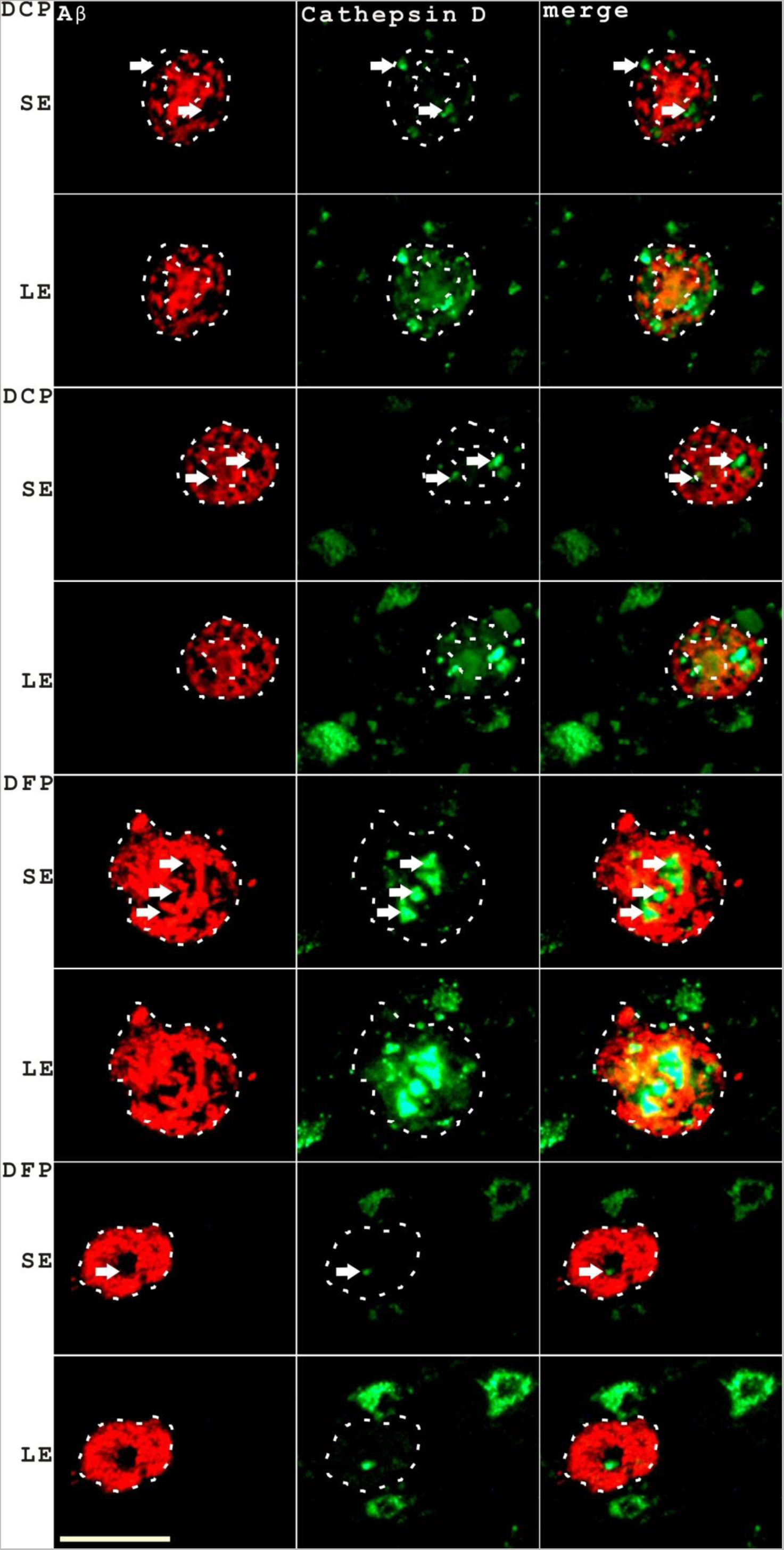
The two distinct layers of Cathepsin D expression in senile plaques were verified with another Aβ/Cathepsin D antibody pair. Dense-core senile plaques showed two layers of Cathepsin D staining using two different exposure settings of the Cathepsin D staining were shown in the top four panels (SE stands for short exposure of 0.5 second while LE stands for long exposure of 1 second). A diffusive senile plaque with two layers of Cathepsin D staining was shown in the fifth and sixth panels. A diffusive senile plaque with very weak plaque Aβ-overlapping Cathepsin D staining was shown in the bottom two panels. The arrows pointed to the strong granule type of Cathepsin D staining while the dashed lines indicated the weaker but constitutive Cathepsin D staining overlapping with senile plaque Aβ staining. Scale bar, 50 μm.

**Supplementary Figure 4.**
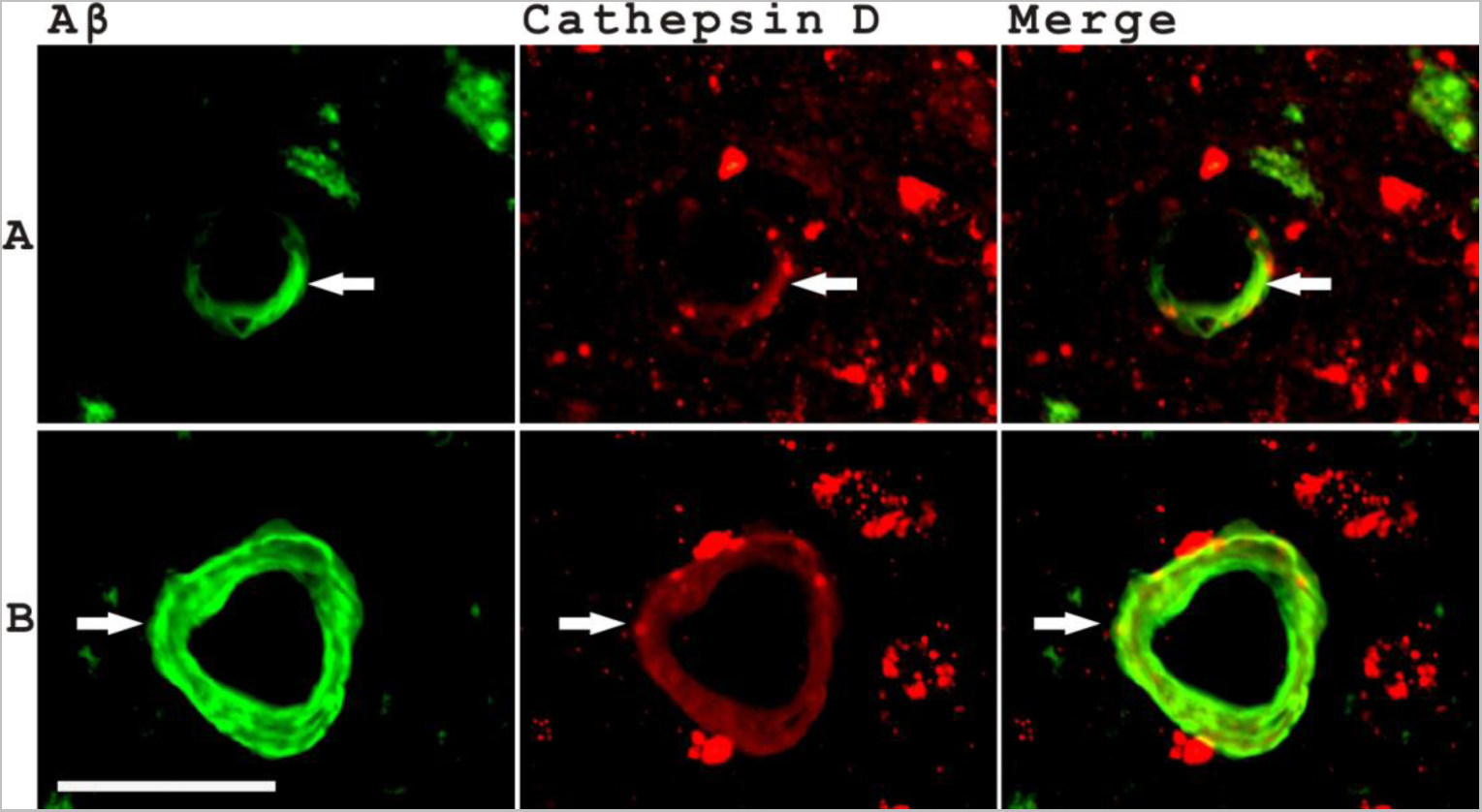
The co-distribution of Aβ and Cathepsin D in CAA. The two panels showed two samples with the co-distribution of Aβ and Cathepsin D in the CAA blood vessels (**A**, **B**). Scale bar, 50μm.

**Supplementary Figure 5.**
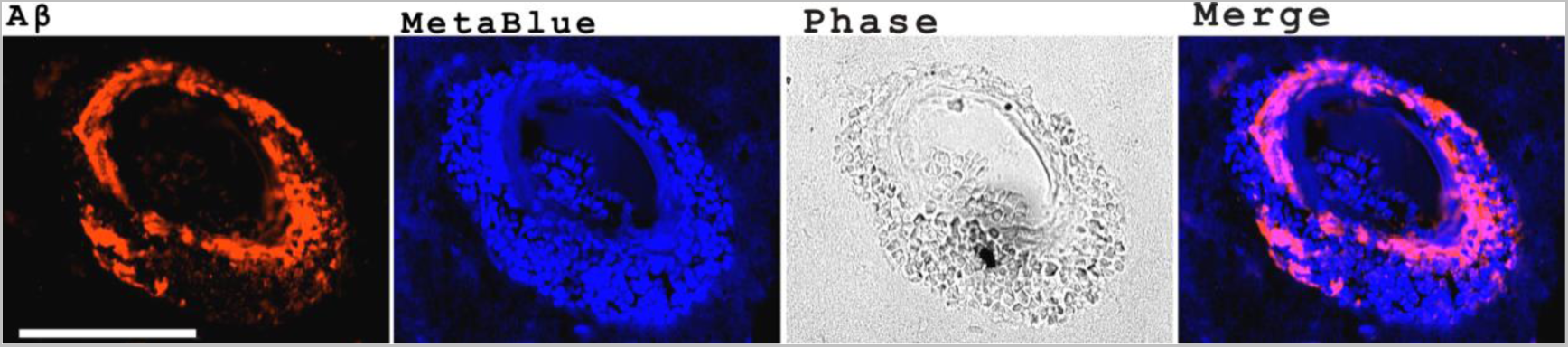
A simple extravagation of red blood cells does not lead to instant senile plaque development. Extravagation of red blood cells out of a CAA blood vessel was observed but with no typical senile plaque structures around. Scale bar, 100μm.

**Supplementary Figure 6.**
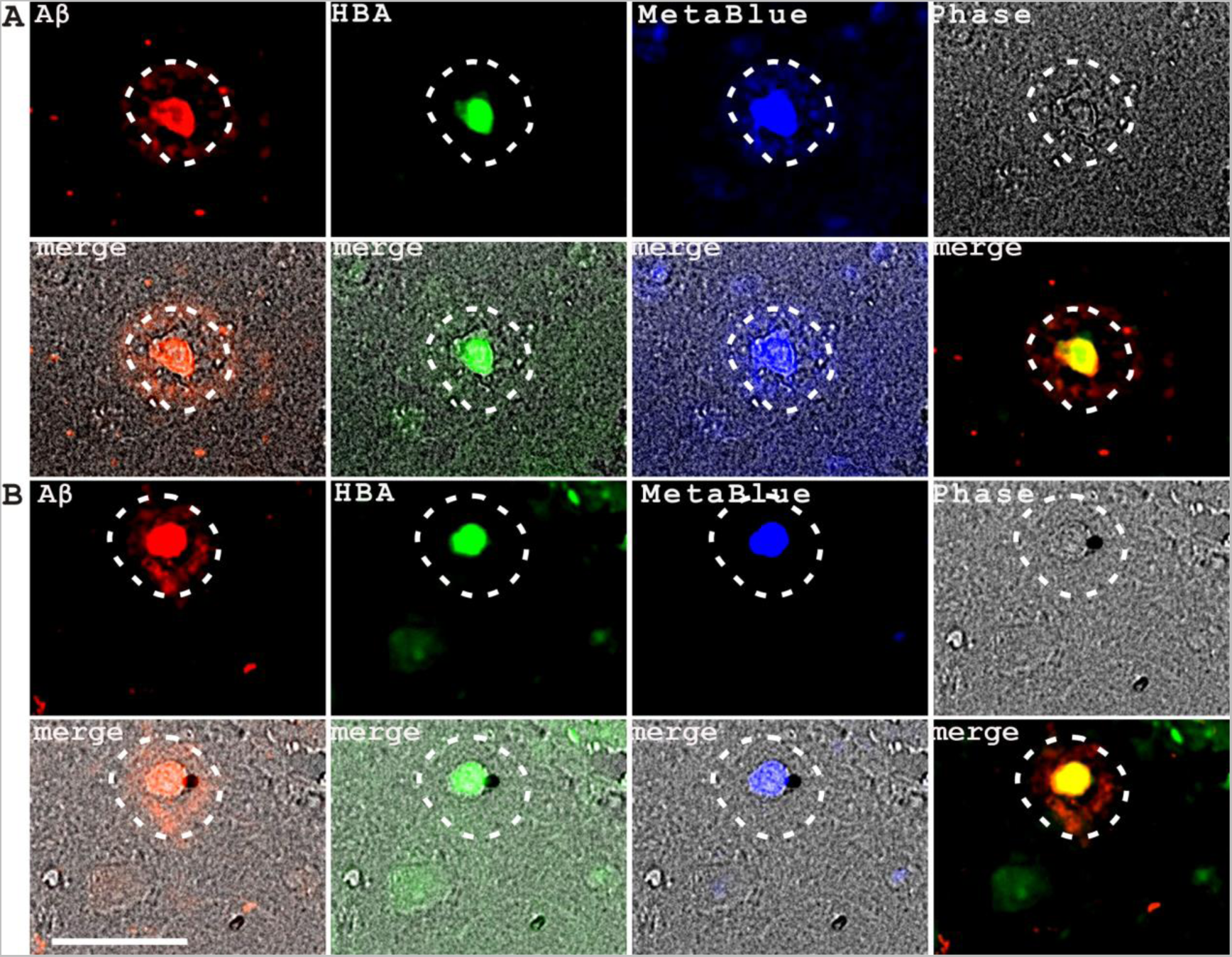
Some dense-core senile plaques had a saccular membrane structure in their dense cores. Aβ and HBA staining of two dense-core senile plaques with also MetaBlue fluorescence and phase contrast showed the existence of saccular membrane structures in the dense cores with concentrated Aβ and HBA and bright MetaBlue signals (**A**, **B**). Scale bar, 50μm.

**Supplementary Figure 7.**
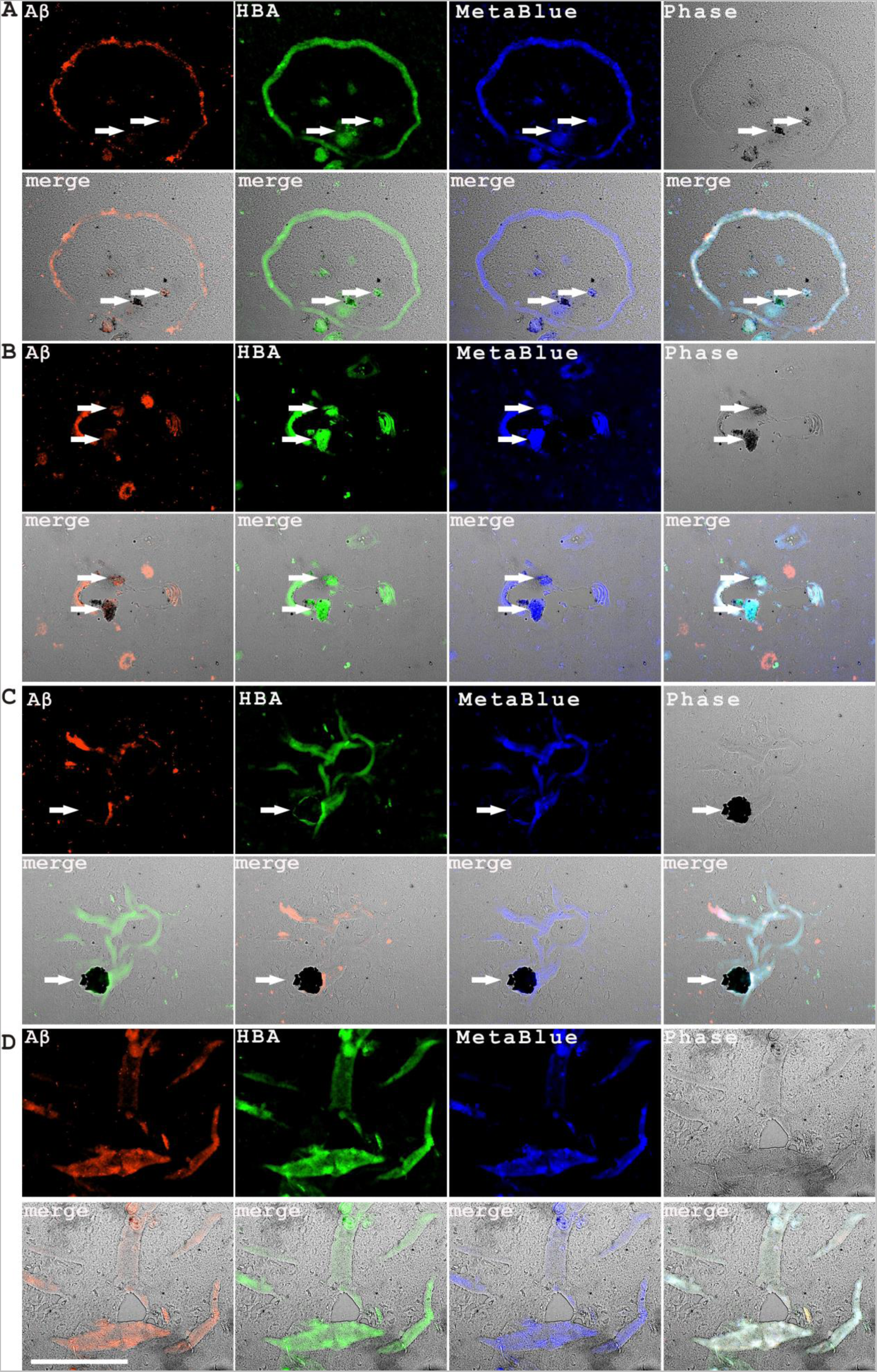
Vascular degeneration in long vessel segments and large vessels or in clustered blood vessels was also observed in AD patient brain tissues. **A.** The top panel showed a long blood vessel fragment with degenerative HBA staining and diffusive Aβ signals but without red blood cells. **B**. The second panel showed a total degeneration of a relatively large vessel with perivascular deposition of HBA and Aβ signals. (**C, D**): The bottom two panels showed simultaneous degeneration of multiple blood vessels clustered together all with diffusive HBA and Aβ staining. To be noticed, black materials that frequently associated with HBA and Aβ immunostaining were often observed around these degenerative tissues (indicated by arrows). Scale bar, 200μm.

**Supplementary Figure 8.**
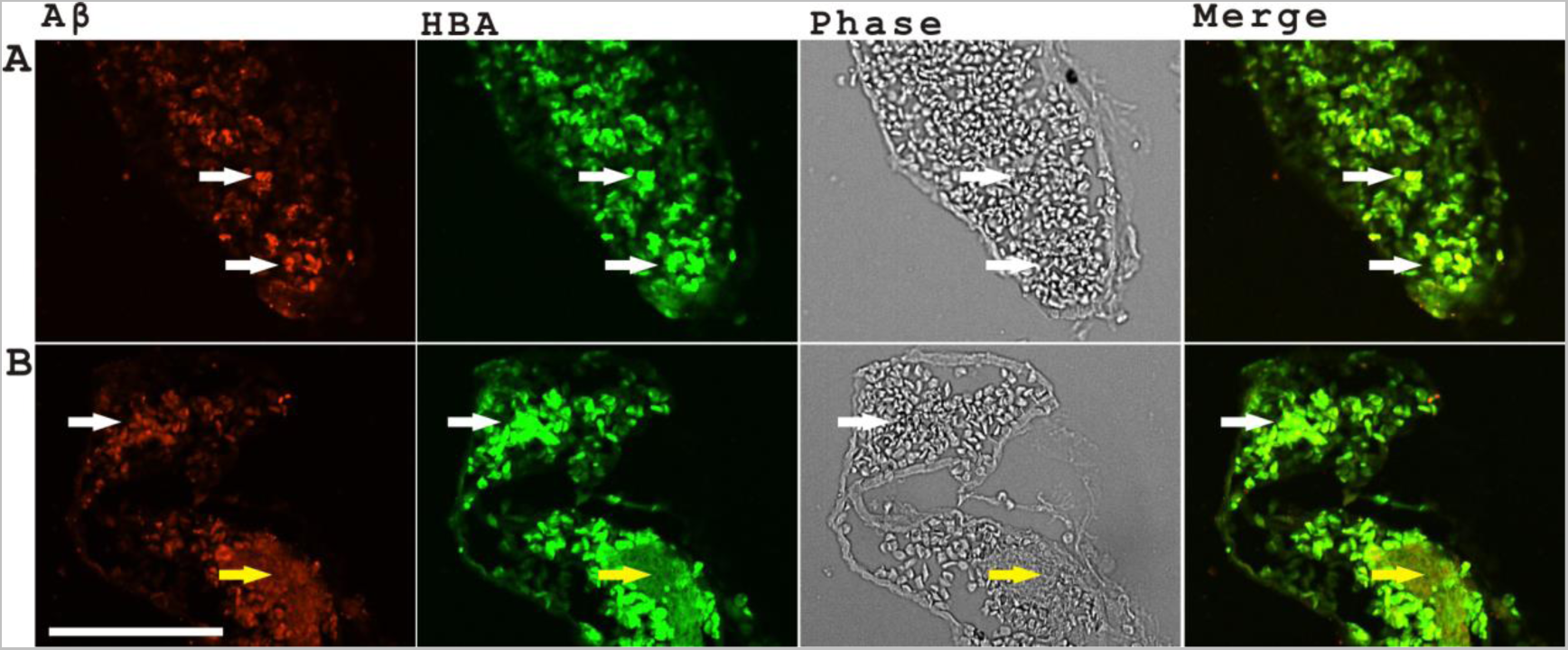
Aβ co-localized with hemoglobin on red blood cells with abnormal morphology resembling schistocytes. **A.** The panel showed many red blood cells with irregular or polygon shapes with the co-localization of Aβ and HBA staining (white arrows). **B**. The panel showed both irregular or polygon shape red blood cells (white arrow) and hemolysis (yellow arrow) with Aβ and HBA co-immunostaining. Scale bar, 100μm.

**Supplementary Figure 9.**
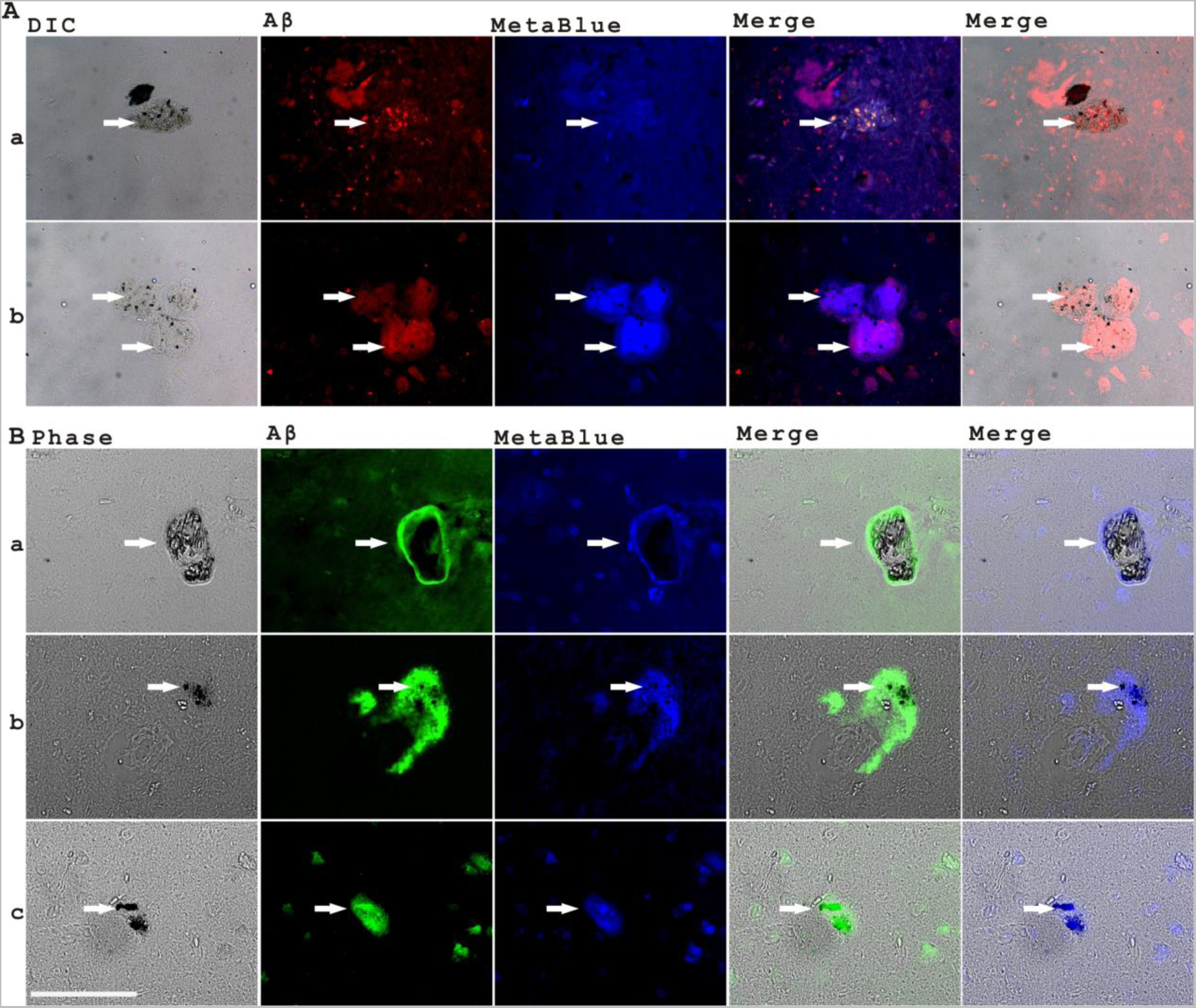
Microaneurysm degeneration was associated with the production of dark-colored materials that could also be detected in some senile plaques. **A.** The two panels showed several saccular microaneurysms containing blue black materials that co-localized with Aβ with red fluorescence and MetaBlue autofluorescence under DIC objective lens (white arrows) (**a, b**). **B**. The top panel showed a saccular microaneurysm containing dark materials under phase contrast objectives that co-localized with Aβ and MetaBlue autofluorescence (white arrows, **a**) while the bottom two panels showed two senile plaques with residual dark materials together with Aβ immunostaining and MetaBlue signals (**b, c**). Scale bars, 100μm.

**Supplementary Figure 10.**
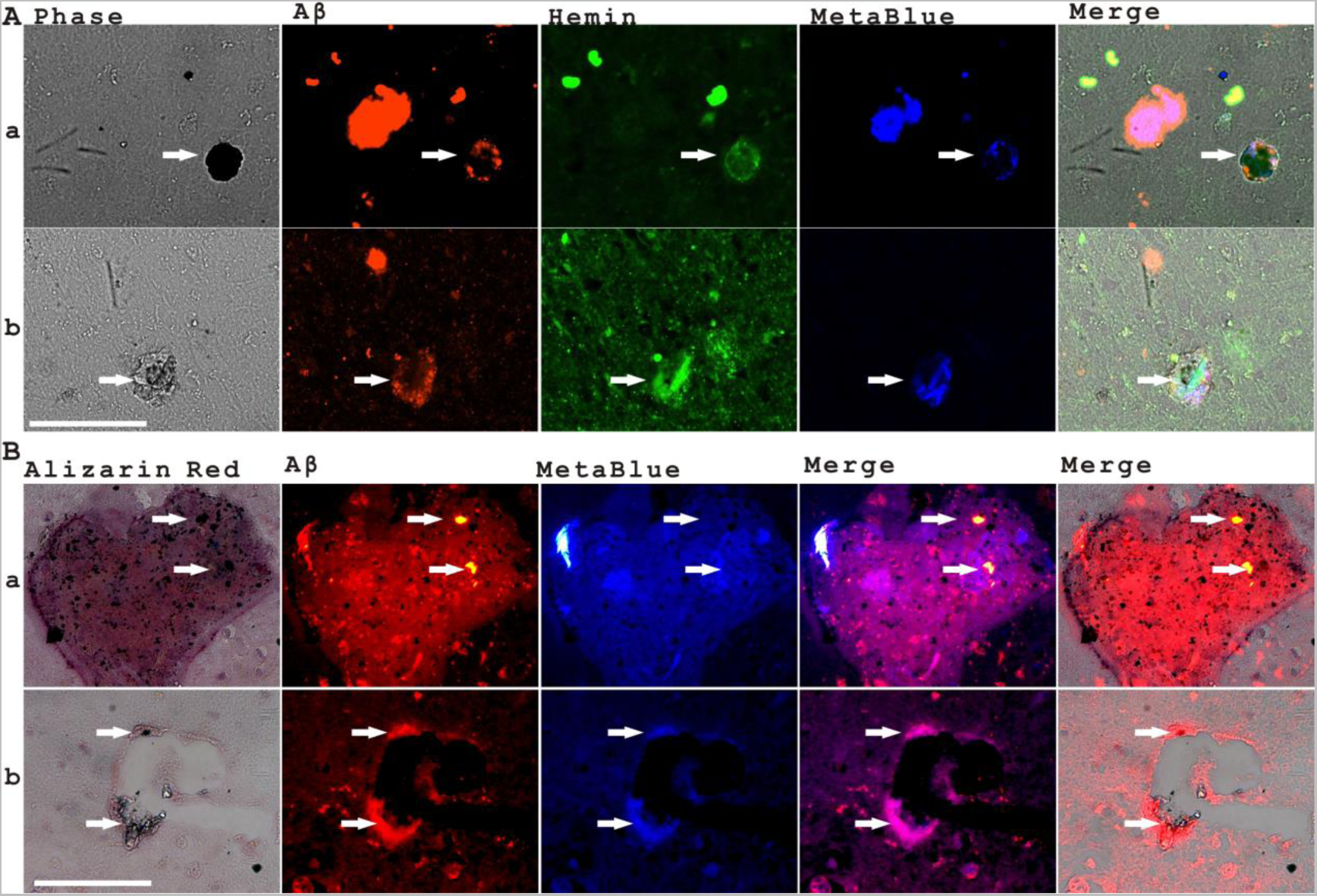
The co-distribution of Aβ and a type of dark materials together with blood markers was observed during microaneurysm formation and vascular degeneration. **A.** The two panels showed several saccular microaneurysm with both Aβ staining and the existence of special dark materials and the co-staining of Hemin and MetaBlue fluorescence (indicated by arrows in **a, b**). **B**. The top panel showed an aneurysm marked with Alizarin Red with the co-distribution of Aβ and dark materials in the aneurysm. The bottom panel showed vascular degeneration with perivascular distribution of Aβ and dark materials. The white arrows indicated the co-distributions of Aβ and dark materials and also MetaBlue fluorescence (indicated by arrows in **a, b**). Scale bars, 100μm.

